# Inositol pyrophosphate profiling reveals regulatory roles of IP6K2-dependent enhanced IP_7_ metabolism in enteric nervous system

**DOI:** 10.1101/2022.09.19.508459

**Authors:** Masatoshi Ito, Natsuko Fujii, Saori Kohara, Shuho Hori, Masayuki Tanaka, Christopher Wittwer, Kenta Kikuchi, Takatoshi Iijima, Yu Kakimoto, Kenichi Hirabayashi, Daisuke Kurotaki, Henning J. Jessen, Adolfo Saiardi, Eiichiro Nagata

**Author notes:** Correspondence: Eiichiro Nagata, M.D., Ph.D., Department of Neurology, Tokai University School of Medicine, 143 Shimokasuya, Isehara, Kanagawa 259-1193, Japan, (ext. 2242), Masatoshi Ito, Ph.D., Support Center for Medical Research and Education, Tokai University, 143 Shimokasuya, Isehara, Kanagawa 259-1193, Japan (ext. 2553).

## Abstract

Inositol pyrophosphates (PP-IPs) regulate diverse physiological processes; to better understand their functional roles, assessing their tissue-specific distribution is important. Here, we profiled PP-IP levels in mammalian organs using a novel HILIC-MS/MS protocol and discovered that the gastrointestinal tract (GIT) contained the highest levels of IP_7_ and its precursor IP_6_. Although their absolute levels in the GIT is diet-dependent, elevated IP_7_ metabolism still exists under dietary regimes devoid of exogenous IP_7_. Of the major GIT cells, enteric neurons selectively express the IP_7_-synthesizing enzyme IP6K2. *IP6K2*-knockout mice exhibited significantly impaired IP_7_ metabolism in the various organs including the proximal GIT. Additionally, HILIC-MS/MS analysis displayed that genetic ablation of *IP6K2* significantly impaired IP_7_ metabolism in the gut and duodenal muscularis externa containing myenteric plexus. Whole transcriptome analysis of duodenal muscularis externa further suggested that IP6K2 inhibition induced the gene sets associated with mature neurons such as inhibitory, GABAergic and dopaminergic neurons, concomitantly with suppression of those for neural progenitor/stem cells and glial cells. In addition, IP6K2 inhibition explicitly affected transcript levels of certain genes modulating neuronal differentiation and functioning, implying critical roles of IP6K2-IP_7_ axis in developmental and functional regulation of enteric nervous system. These results collectively reveal an unexpected role of mammalian IP_7_—a highly active IP6K2-IP_7_ pathway is conducive to enteric nervous system.

## Introduction

Myo-inositol phosphates (IPs) are ubiquitously synthesized in all organisms and are involved in pleiotropic biological processes, most importantly in intracellular signaling (Irvine and Schell, 2001). Among the IP family, inositol hexakisphosphate (IP_6_) is the most abundant, and serves as a precursor of inositol pyrophosphates (PP-IPs) possessing diphosphate moieties at specific carbon positions (Saiardi, 2012; Wilson et al, 2013; Shears, 2015; Shah et al, 2017). Diphosphoinositol pentakisphosphate (IP_7_) and bisdiphosphoinositol tetrakisphosphate (IP_8_) are the most well-characterized PP-IPs in mammals and yeasts, and carry diphosphate moieties at the 5-position (5-IP_7_) and 1,5-positions (1,5-IP_8_) of the inositol ring, respectively (Draskovic et al, 2008; Shears, 2015). Recent studies using mammalian cells have demonstrated that PP-IPs regulate phosphate flux, energy homeostasis, and post-transcriptional processes at the molecular level (Wilson et al, 2019; Li et al, 2020; López-Sánchez et al, 2020; Sahu et al, 2020; Gu et al, 2021). In mammals, 5-IP_7_ is synthesized by three inositol hexakisphosphate kinases (IP6Ks) IP6K1, IP6K2, and IP6K3. IP6K1 and IP6K2 are expressed in most mammalian tissues, with highest expression in the brain and testis, whereas IP6K3 expression is mainly restricted to the muscles (Saiardi et al, 2001; Moritoh et al, 2016; Laha et al, 2021). *In vivo* studies using *IP6K1*- or *IP6K2*-knockout mice suggest that PP-IPs contribute to the development and maintenance of neuronal cells (Fu et al, 2017; Nagpal et al, 2018, 2021). In addition to these *in vivo* mice studies, our as well as other research groups have shown that PP-IPs are pathophysiologically involved in the progression of obesity (Chakraborty et al, 2010; Ghoshal et al, 2016), in cancer (Rao et al, 2015) and in neurodegenerative disorders such as Huntington’s disease (Nagata et al, 2011), amyotrophic lateral sclerosis (Nagata et al, 2016), and Alzheimer’s disease (Crocco et al, 2016). Therefore, PP-IPs are currently being considered as potential therapeutic targets for several diverse human disorders (Shears, 2016; Chakraborty, 2018). However, we are unaware of any systematic studies that have directly and comprehensively analyzed PP-IP distribution in mammalian tissues, which could provide valuable insights into the effects of pharmacological interventions on the PP-IP system.

Over the past decade, extensive efforts have been made to develop analytical methods for detecting PP-IPs. Traditionally, PP-IPs have been studied using radioisotopic ^3^H-inositol labeling coupled with anion exchange chromatography (Azevedo and Saiardi, 2006), which allows sensitive detection of metabolically labeled PP-IPs from cultured cells. Electrophoretic separation and colorimetric visualization of PP-IPs (Losito et al, 2009) have also become alternative standard methods for distinguishing PP-IPs. However, PP-IPs in mammalian tissues can neither be radioisotopically labelled, nor explicitly detected using colorimetric visualization. A mass spectrometric method coupled with capillary electrophoretic separation (capillary electrophoresis-mass spectrometry, CE-MS) (Qiu et al, 2020) was recently reported for sensitive analysis of PP-IPs in biological samples at the isomer level. However, the instrument setup involved is complex and requires skillful handling, and is therefore rarely available in research institutes.

We recently developed an analytical method that directly detects mammalian-derived IP_7_ and its precursor IP_6_ using conventional liquid chromatography-tandem mass spectrometry (LC-MS/MS) coupled with hydrophilic interaction liquid chromatography (HILIC) (Ito et al, 2018), enabling the previously impossible quantitation of PP-IPs in mammalian tissues. In this study, we analyzed PP-IP and their precursor IP_6_ levels in mammalian organs using a refined HILIC-MS/MS protocol. We found that IP_7_ was present at explicit levels in the mammalian central nervous system (CNS), where IP6Ks are highly expressed. Surprisingly, we also discovered that the highest IP_7_ production was observed in the gastrointestinal tract (GIT), even after depletion of dietary derived IP_7_. Of the major GIT cells, enteric neurons selectively expressed IP_7_-synthesizing enzyme IP6K2, which was revealed by assessment of single cell RNA-sequencing (scRNA-seq) data sets and confirmed by immunohistochemical detection. Our HILIC-MS/MS survey using *IP6K2*-knockout (*IP6K2*^-/-^) mice exhibited that IP6K2-dependent enhanced IP_7_ metabolism exists in the gut and duodenal muscularis externa where myenteric plexus is located. We further performed whole transcriptome analysis of *IP6K2*-deficient and wild type (WT) duodenal muscularis externa to define a physiological role of IP6K2-IP_7_ pathway in enteric nervous system (ENS).

## Results

### Refinement of HILIC-MS/MS protocol for PP-IP analysis

Before investigating PP-IP metabolism in mammalian tissues, we improved our HILIC-MS/MS analysis protocol for unequivocal detection and more precise quantitation of PP-IPs. Medronic acid compatible with LC-MS analysis significantly improves the chromatographic peak shape of phosphorylated compounds (Hsiao et al, 2018). We employed a form of this solvent additive that has been optimized for HILIC analysis (InfinityLab deactivator additive, Agilent Technologies) and found that using it significantly improved the peak shapes of IP_6_ and IP_7_ (Fig. 1A), whereas without the additive, remarkably poor IP_6_ and IP_7_ peak shapes were obtained, probably due to the cumulative adsorption of cationic contaminants on the column derivative (amino group). Chromatographic peaks of IP_7_ levels as low as 10 pmol were discernible using this additive (Fig. 1B). Direct use of LC-MS grade medronic acid resulted in similar beneficial effects on IP_6_ and IP_7_ peaks (Fig. S1), but the background noise was relatively higher than that for the InfinityLab deactivator additive. Thus, the additive was used in all our subsequent analyses.

**Figure 1.**
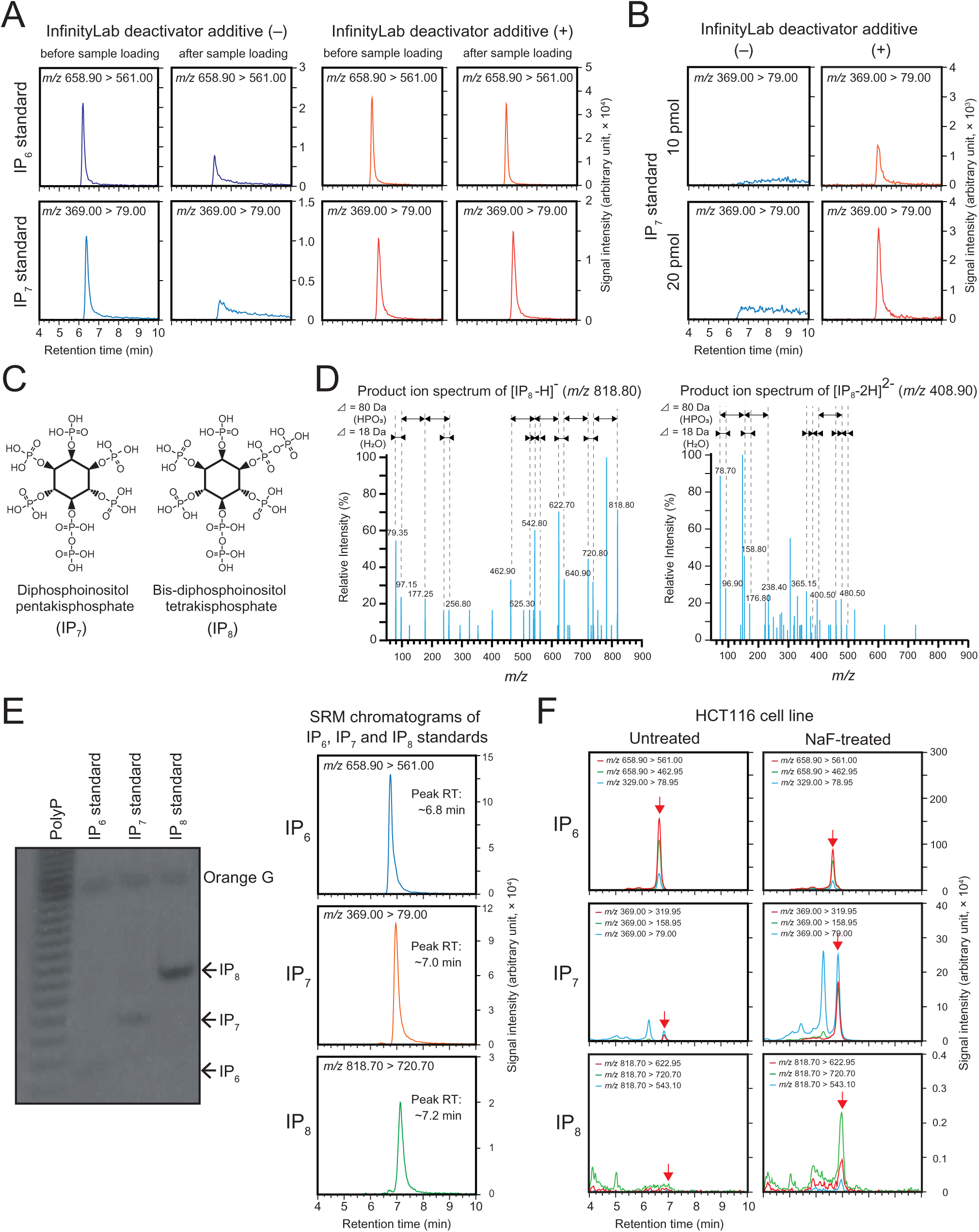
Refined HILIC-MS/MS analysis for IP_6_ and PP-IPs. **A.** Effect of InfinityLab deactivator additive as a mobile phase modifier on SRM chromatograms of IP_6_ and IP_7_ before and after biological sample injection. 100 pmol of each synthetic analyte were injected. **B.** Effect of InfinityLab deactivator additive as a mobile phase modifier on the detection of low amounts of synthetic PP-IP. 10 and 20 pmol of IP_7_ standard were injected. **C.** Chemical structure of IP_7_ and IP_8_. **D.** Product ion spectrum of IP_8_ (singly deprotonated precursor, left panel; doubly deprotonated precursor, right panel). Characteristic fragment ions generated by loss of water (H_2_O, 18□Da) and phosphoric acid (H_3_PO_4_, 80□Da) are also shown. **E.** Gel electrophoretic results (left panel) and SRM chromatograms (right panel) of synthetic IP_6_, IP_7_, and IP_8_ standard. The PolyP ladder was used as an electrophoresis standard. 500 pmol of each standard were injected for LC-MS. **F.** Representative SRM chromatograms of IP_6_, IP_7_, and IP_8_ in untreated (left panel) and NaF-treated (right panel) HCT116 cell samples. The three best transitions per molecule are shown for the peak identification of each compound. Arrows indicate the SRM peak of corresponding analytes.

To quantitate IP_8_ (Fig. 1C) simultaneously with IP_6_ and IP_7_, we assessed the mass spectra of IP_8_ fragment ions obtained by collision-induced dissociation of the synthetic IP_8_ standard (Fig. 1D). A series of fragment ions representing the losses of phosphate (80 Da) and water (18 Da) appeared in the spectra. Based on this result, we assigned each IP_8_ fragment and optimized the selected reaction monitoring (SRM) conditions (Table S1). Using chemical standards of PP-IPs and their precursor IP_6_, we observed chromatographic peaks of IP_6_, IP_7_, and IP_8_ at regular intervals of 0.2 min (Fig. 1E). To benchmark this method for the detection of endogenous PP-IPs, we treated HCT116 cells with NaF, which is known to increase IP_7_ level (Menniti et al, 1993). While a clear IP_6_ SRM peak and subtle IP_7_ and IP_8_ SRM peaks were observed for untreated HCT116 cells, explicit IP_7_ and IP_8_ SRM peaks were detected for NaF-treated cells (Fig. 1F). We also observed a dose-dependent reduction in IP_7_ level and the IP_7_/IP_6_ ratio in HCT116 cells treated with the IP6K inhibitor TNP (Fig. S2). Thus, our refined HILIC-MS/MS protocol achieved robust, sensitive, and reliable detection of endogenous IP_6_, IP_7_, and IP_8_ in biological samples.

### The mammalian gastrointestinal tract (GIT) contains high levels of PP-IPs

Using the newly developed HILIC-MS/MS protocol, we investigated the distribution of PP-IPs in experimental model rodents fed with a standard plant-based diet (CE-2; Clea, Japan). Fifteen organs, including the CNS and GIT, were harvested from standard diet-fed C57BL/6J male mice. Surprisingly, HILIC-MS/MS analysis showed that the GIT had the highest levels of IP_6_ and IP_7_, even after extensive rinsing of the organs with phosphate-buffered saline (PBS) to wash out the digested contents (Fig. 2A, B and Table S2). Importantly, the IP_7_/IP_6_ ratio in the GIT was remarkably high, by far the highest in all organs examined (Fig. 2C and Table S2). A subtle IP_8_ SRM peak was detected in stomach and small intestine samples, wherein IP_7_ was abundant (Fig. 2D) but was not detected in other organs. While IP_7_ SRM peaks were clearly detected in CNS samples (Fig. 2E), IP_7_ levels in the CNS were modest compared with those in the GIT. Moreover, the IP_7_/IP_6_ ratio in the spinal cord appeared to be higher than that in the cerebrum (Fig. 2C).

**Figure 2.**
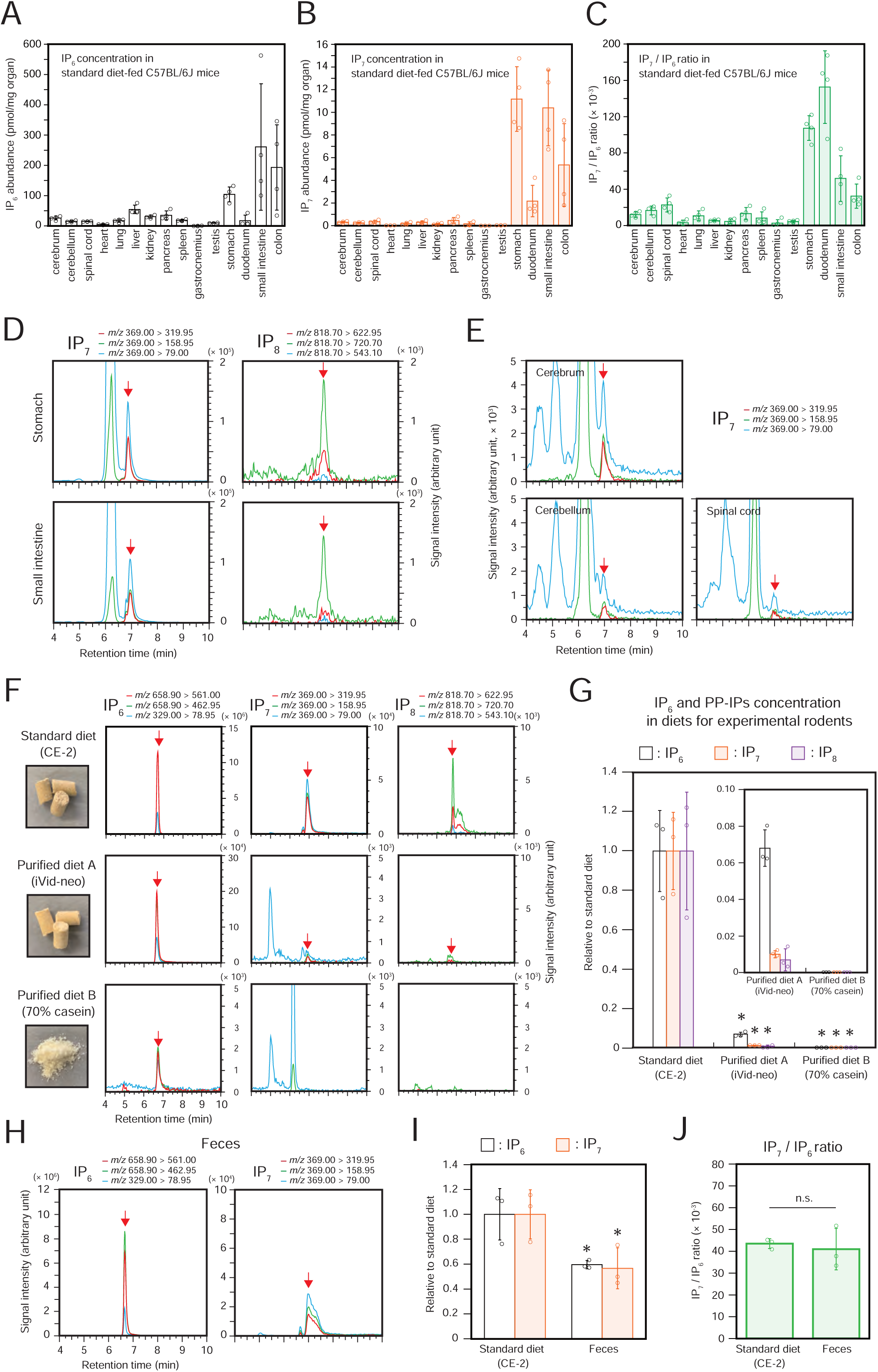
The mammalian gastrointestinal tract (GIT) contains high levels of PP-IPs. **A-C.** The concentrations of IP_6_ **(A)** and IP_7_ **(B)** and IP_7_/IP_6_ ratios **(C)** in the 15 organs of standard diet-fed C57BL/6J mice. The values shown are expressed as pmol per mg of organ weight (n = 4). **D.** Representative SRM chromatograms of IP_7_ and IP_8_ in stomach and small intestine samples of standard diet-fed C57BL/6J mice. The three best transitions per molecule are shown for the peak identification of each compound. The arrows indicate the SRM peaks of the corresponding analytes. **E.** Representative SRM chromatograms of IP_7_ in CNS samples of standard diet-fed C57BL/6J mice. **F.** Photographs (left) and SRM chromatograms of IP_6_, IP_7_, and IP_8_ (right) of standard diet (CE-2) and the two different purified diets (iVid-neo and 70% casein). **G.** Relative concentrations of IP_6_ and PP-IPs in the standard and purified diets. The values shown represent the mean□±□standard deviation (SD) of three independent experiments and are expressed relative to those of the standard diet. Asterisks indicate statistical significance (P□<□0.05, Student’s t-test) compared with the standard diet. **H.** Representative SRM chromatograms of IP_6_ and IP_7_ in the feces of standard diet-fed C57BL/6J mice. **I.** Relative concentrations of IP_6_ and IP_7_ in the feces of standard diet-fed C57BL/6J mice. The values shown are expressed relative to those of the standard diet (n = 3). Asterisks indicate statistical significance (P□<□0.05, Student’s t-test) compared with the standard diet. **J.** IP_7_/IP_6_ ratio in standard diet and feces of standard diet-fed C57BL/6J mice (n = 3). n.s., not significant (Student’s *t*-test).

Several reports have shown that IPs (mainly IP_6_, known as phytic acid) are present in a variety of crop seeds (Dorsch et al, 2003; Liu et al, 2009; Kolozsvari et al, 2015; Duong et al, 2017); moreover, plants also generate PP-IPs, which are crucial for phosphorus-starvation responses (Dong et al, 2019; Ried et al, 2021; Riemer et al, 2021). Therefore, we assumed that the plant-based CE-2 diet contains IP_6_ and PP-IPs, and explicit chromatographic peaks of IP_6_, IP_7_, and IP_8_ were observed in CE-2 samples (Fig. 2F, upper panels). The concentrations of IP_6_ and IP_7_ in CE-2 were 3.96 ± 0.82 nmol/mg and 0.17 ± 0.03 nmol/mg, respectively. We next investigated their concentrations in purified diets with minimal levels of plant-derived components (Fig. 2F, middle and lower panels). The two purified diets examined (iVid-neo and 70% casein) contained low amounts of IP_6_ and negligible amounts of IP_7_ and IP_8_. Quantitative analysis revealed that the levels of all PP-IPs in both purified diets were less than 2% of those in CE-2 (Fig. 2G).

Since the IP_7_/IP_6_ ratios in the stomach and duodenum were significantly higher than that in CE-2 (Fig. 2C), the high IP_7_ level detected could not be attributed to its direct absorption from CE-2 diet, so it must have been endogenously produced by active IP6K enzyme. However, to exclude the possibility of selective intestinal absorption of IP_7_, we analyzed the feces of mice fed on CE-2 and estimated the loss of IP_6_ and IP_7_ in the digestive system (Fig. 2H). Similar to those for CE-2 samples, IP_6_ and IP_7_ SRM peaks were clearly observed in mouse feces samples. Quantitative analysis showed that approximately 50% of IP_6_ and IP_7_ in ingested food remained in the feces (Fig. 2I). Since the IP_7_/IP_6_ ratio remained unchanged between undigested CE-2 and feces, we could exclude that IP_7_ is selectively absorbed in the GIT, further demonstrating that the abundant IP_7_ levels observed in the GIT must be endogenously generated by cellular metabolism.

### Enhanced IP_7_ metabolism is retained in the proximal GIT of rodents under conditions of depleted dietary IP_6_ and PP-IP supply

To validate the presence of endogenously synthesized PP-IPs in the GIT, C57BL/6J mice were fed for 2 months on standard CE-2 diet; on iVid-neo containing negligible amounts of IP_6_ and PP-IPs (Fig. 2E); or left fasting for 48 h (Fig. 3A). The GIT of both purified diet-fed and fasted mice showed a reduction in IP_6_ and IP_7_ levels compared with those of standard diet-fed mice; however, the levels were still close to (in the case of IP_6_) or far greater than (in the case of IP_7_) the CNS levels (Fig. 3B and C). IP_8_ was not detected in any of the tested organs of purified diet-fed and fasted mice. The SRM chromatograms of both purified diet-fed and fasted mice samples had explicit IP_7_ SRM peaks (Fig. 3D). Importantly, the stomach and duodenum of these mice showed prominently higher IP_7_/IP_6_ ratio than those of their standard diet-fed counterparts (Fig. 3E), implying further enhanced IP_7_ metabolism compensated for the overall reduced IP_7_ level. On the other hand, both purified diet-fed and fasted mice did not show any changes in the IP_6_ and IP_7_ levels as well as the IP_7_/IP_6_ ratio in the CNS and testis compared with those of mice fed a standard diet. However, as with standard diet-fed mice, both purified diet-fed and fasted mice showed higher IP_7_/IP_6_ ratios in the spinal cord than in the cerebrum. We also investigated IP_7_ levels in the GIT of purified diet (70% casein)-fed Sprague–Dawley rats (Fig. 3F). Analogous to the results observed in the mouse model, both IP_6_ and IP_7_ levels in the GIT of these rats were drastically reduced compared with those in the standard diet-fed GIT and comparable to those in the CNS (Fig. 3G and H). In addition, the IP_7_/IP_6_ ratio was higher in the stomach and duodenum of purified diet-fed rats compared with that of standard diet-fed rats (Fig. 3I), further demonstrating very active IP_7_ metabolism in the mammalian proximal GIT.

**Figure 3.**
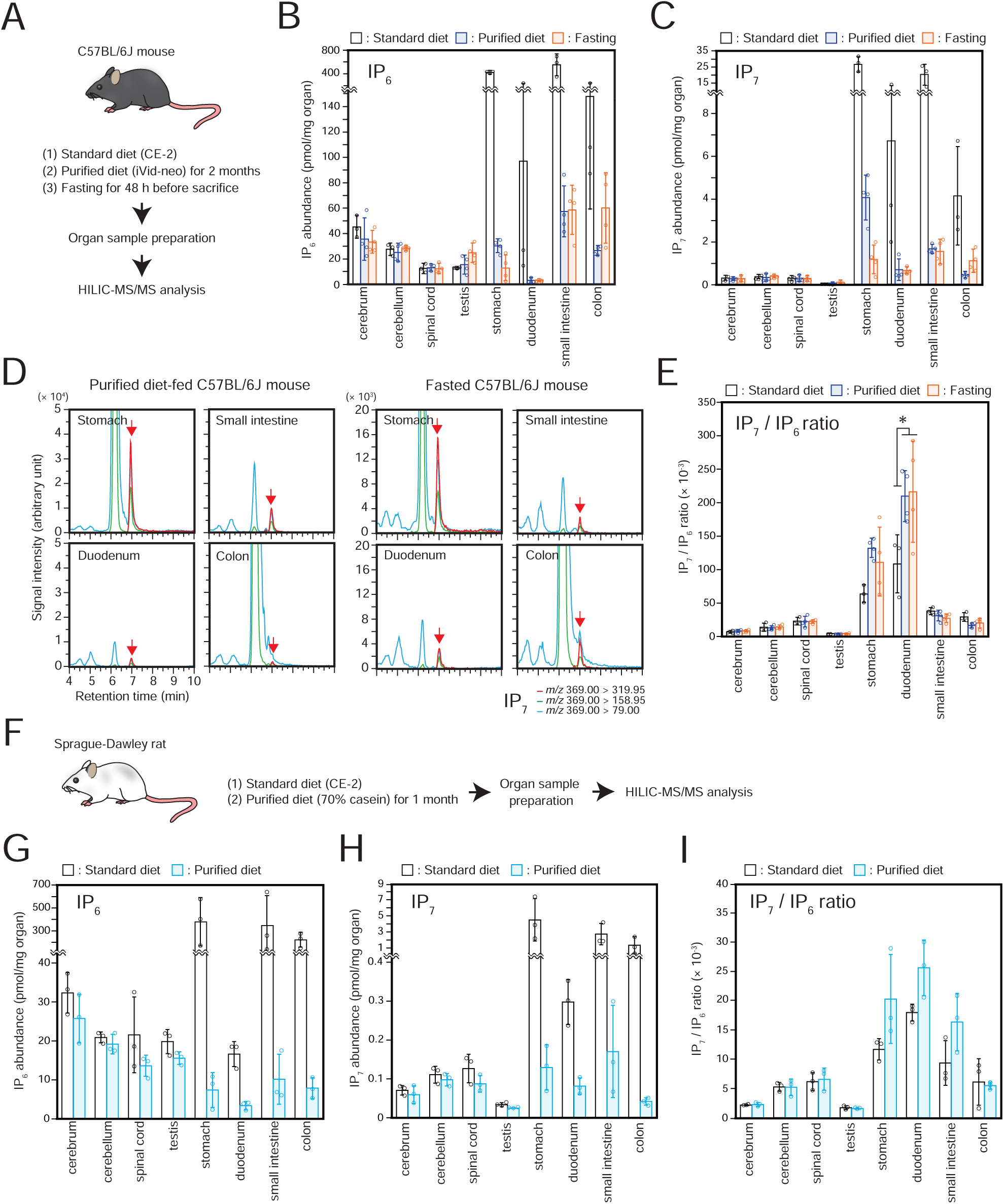
Enhanced IP_7_ metabolism is retained in the proximal GIT of rodents under conditions of depleted dietary IP_6_ and PP-IP supply. **A.** Schematic illustration of the experimental workflow. C57BL/6J mice were fed a standard diet (n = 3) or purified diet (iVid-neo) for 2 months (n = 4) or fasted for 48 h (n = 4). **B, C.** The concentrations of IP_6_ **(B)** and IP_7_ **(C)** in the CNS, testes, and GIT of C57BL/6J mice under the three different conditions. The values shown are expressed as pmol per mg of organ weight. **D.** Representative SRM chromatograms of IP_7_ in the GIT of C57BL/6J mice fed with purified diet (left panel) or under fasting conditions (right panel). Arrows indicate the SRM peak of IP_7_. **E.** IP_7_/IP_6_ ratios in the CNS, testes, and GIT of C57BL/6J mice under the three different conditions. Asterisk indicates statistical significance (P□<□0.05, one-way ANOVA, Bonferroni-type post-hoc test) compared with the standard diet-fed mice. **F.** Graphical scheme of the experiment. Sprague–Dawley rats were fed a standard (n = 3) or purified diet (70% casein) for 1 month (n = 3). **G-I.** Concentrations of IP_6_ **(G)**, IP_7_ **(H)** and IP_7_/IP_6_ ratios **(I)** in the CNS, testes, and GIT of the rats under the two different conditions. The values shown are expressed as pmol per mg of organ weight.

### Enteric neurons highly express IP6K2 in the mammalian GIT

To investigate the expression levels of the three IP6Ks in each GIT cell type, we used single-cell RNA sequencing (scRNA-seq) datasets and compared the expression levels of IP6Ks among GIT cell types. Quantitative analysis using a human embryonic intestinal cell scRNA-seq dataset (Fawkner-Corbett et al, 2021) showed that enteric neural cells expressed the highest levels of IP6K2 among different intestinal cells (Fig. 4A, left panel). In enteric neural cells, IP6K2 was selectively expressed across enteric neuron subsets, such as motor neurons, interneurons, and neuroendocrine cells, but not in glial cells (Fig. 4A, right panel). This analysis was further supported by the IP6K quantitation using both E15.5 (Fig. S3A) and E18.5 (Fig. 4B) mouse embryonic ENS scRNA-seq datasets (Morarach et al, 2021). As in humans, IP6K2 isoform expression level in mouse enteric neurons was higher than in other neural cells such as neuroblasts, progenitors, glial cells, and Schwann cells. Moreover, the transcriptional analysis-based data were verified using immunohistochemical analyses. IP6K2 colocalized with the neuronal marker HuC/D in the mouse duodenal muscle layer, suggesting IP6K2 was expressed in the myenteric plexus (Fig. 4C). Other than enteric neurons, several cell types, including secretory progenitor cells, also expressed relatively high levels of IP6K2 (Fig. S3B). In addition, mouse enteric epithelial cell scRNA-seq data (Haber et al, 2017) showed that IP6K2 is expressed in mouse enteroendocrine cells (Fig. S3C). Expression levels of IP6K1 and IP6K3 in entire embryonic intestinal cells were low and negligible, respectively (Fig. 4A and B). These results suggest that IP6K2 is highly expressed in mammalian enteric neurons.

**Figure 4.**
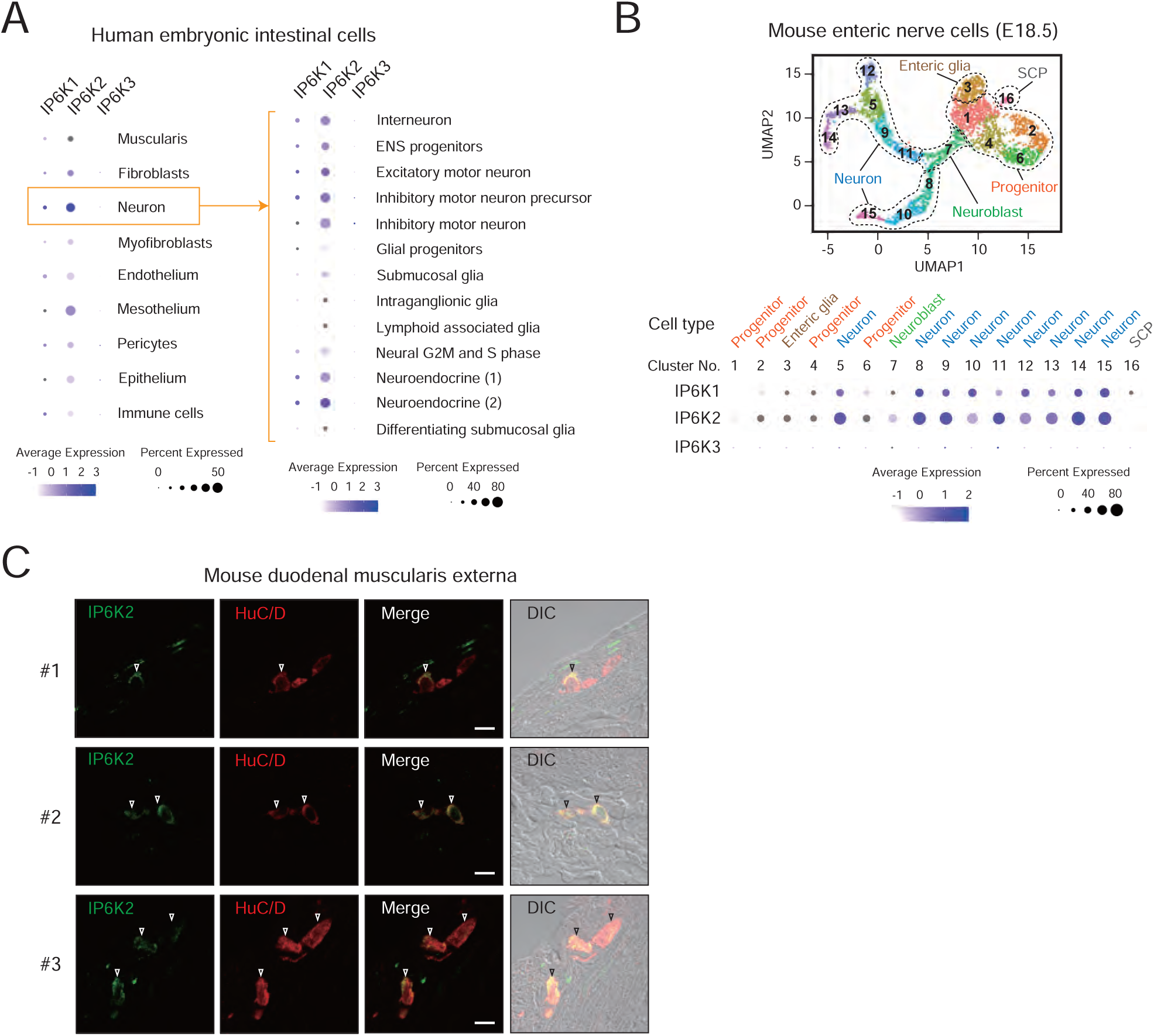
Enteric neurons highly express IP6K2 in the mammalian GIT. **A.** Expression analysis of IP6K1-3 in intestinal cell subsets using publicly available scRNA-seq datasets. Relative expression (log scale) of IP6K1-3 among human embryonic enteric cells (left) and their neural cell subsets (right), obtained by analysis of human embryonic intestinal cells scRNA-seq datasets are shown (Fawkner-Corbett et al, 2021). The size and color of the dots represent the percentage of cells which express IP6K1-3 mRNA and their average abundances within a cluster, respectively. **B.** UMAP-based unsupervised clustering of recently reported mouse embryonic (E18.5) ENS data (Morarach et al, 2021) (upper panel). Assignment of cell identities was based on the expression of signature genes as described in the literature: *Sox10* (Progenitor), *Ascl1* (Neuroblast), *Elavl4* (Neuron), *Plp1* (Enteric glia) and *Dhh* (SCP). Relative expression (log scale) of IP6K1-3 among the ENS clusters (lower panel) are shown. ENS, enteric nervous system; SCP, Schwann cell precursor; E, embryonic day; UMAP, uniform manifold approximation and projection. **C.** Immunohistochemical analysis of IP6K2 expression in the duodenal muscularis externa of C57BL/6J mice. Three different areas of confocal microscopy images are shown. The neuronal marker HuC/D was also detected to identify enteric neurons in the myenteric plexuses. Open arrowheads indicate double-positive cells. DIC images were overlaid onto the respective merged fluorescent images to identify cell contours. DIC, differential interference contrast. Scale bar = 10 μm.

### *IP6K2*^-/-^ mice show significant impairment of IP_7_ metabolism in the proximal GIT

To estimate the importance of IP6K2 in endogenous IP_7_ synthesis in the mammalian organs including GIT, we employed a genetically modified mouse in which *IP6K2* exon 6, encoding the kinase domain, was specifically deleted (Fig. 5A, upper panel) (Rao et al, 2014). To avoid any contamination of dietary-derived IPs in our analysis, *IP6K2*-knockout (*IP6K2*^-/-^) or wild type (WT) mice raised on the standard CE-2 diet were switched to a purified diet (iVid-neo) for one week, and then fasted for 48 h before sacrifice (Fig. 5A, lower panel). In WT mice, IP6K2 mRNA containing the exon 6 sequence was expressed in the proximal GIT but only marginally compared with the expression in the CNS (Fig. 5B). As expected, the IP6K2 transcript was absent in *IP6K2*^-/-^ mouse organs. We confirmed the loss of IP6K2 expression at the protein level using cerebrum lysate (Fig. 5C), because it has high IP6K2 protein expression and thus was useful for clearly validating the loss of IP6K2 in *IP6K2*^-/-^ mice. HILIC-MS/MS analysis showed that *IP6K2*^-/-^ mice had significantly lower levels of IP_7_ in various organs, including the stomach and duodenum, compared with those in their WT counterparts, while IP_6_ levels in each organ were almost the same between *IP6K2*^-/-^ and WT mice (Fig. 5D and E). As previously observed (Fig. 3E and I), the IP_7_/IP_6_ ratios in the stomach and duodenum of WT mice were much higher than those in the other organs examined (Fig. 5F). On the other hand, the IP_7_/IP_6_ ratios in these two organs were significantly reduced in *IP6K2*^-/-^ mice. The IP_7_ SRM peaks for the *IP6K2*^-/-^ mouse stomach and duodenum were also smaller compared with those of WT mice, while IP_6_ levels were unchanged (Fig. 5G and H). Collectively, these data demonstrate that IP6K2 is required for enhanced IP_7_ metabolism in the mammalian proximal GIT.

**Figure 5.**
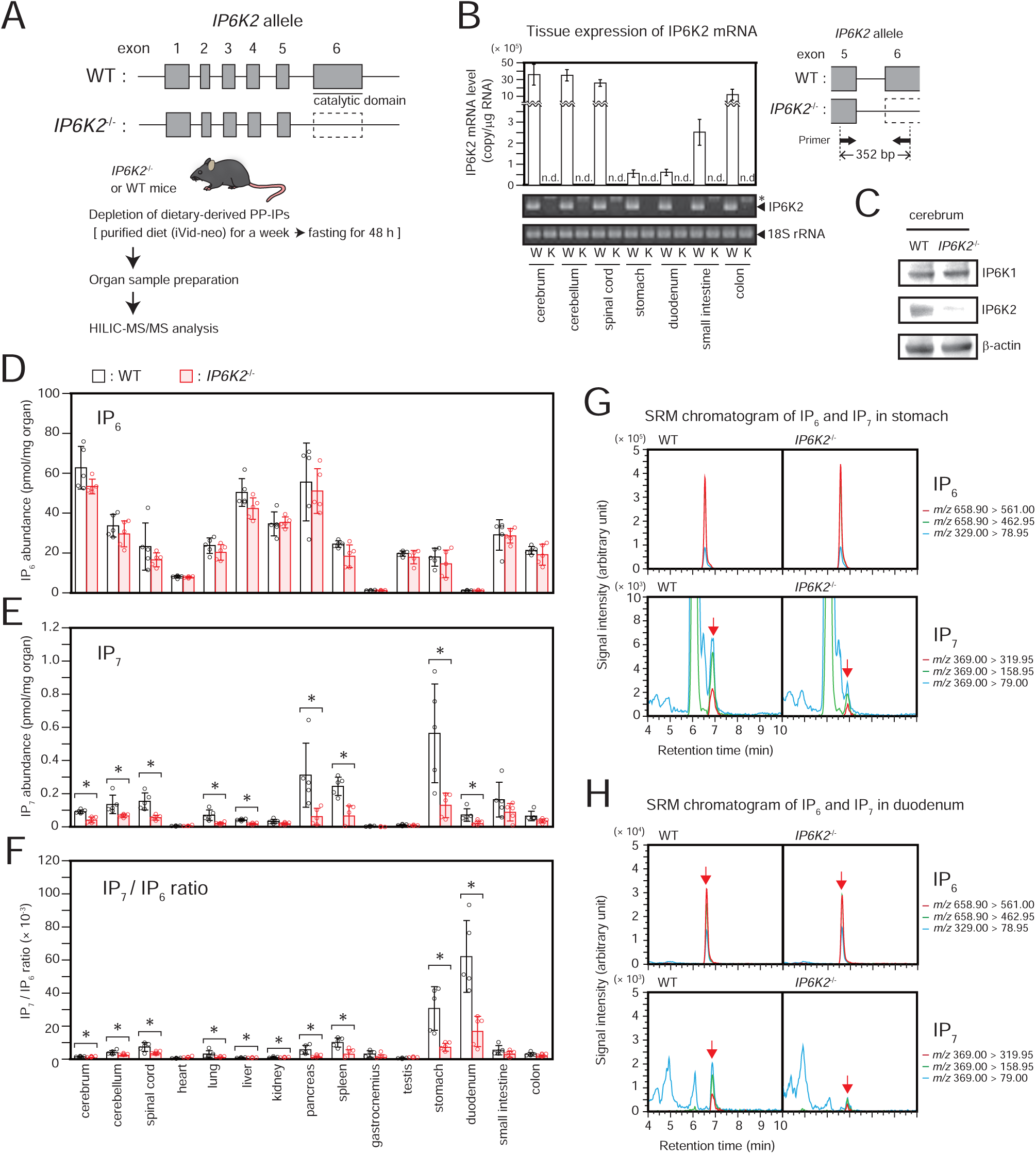
*IP6K2*^-/-^ mice show significant impairment of IP_7_ metabolism in the proximal GIT. **A.** Schematic depiction of the *IP6K2* genomic locus in *IP6K2*^-/-^ and WT mice (upper panel) and the experimental workflow (lower panel). IP6K2 exons and introns are represented as boxes and lines, respectively. **B.** IP6K2 mRNA levels in the CNS and GIT of *IP6K2*^-/-^ and WT mice (upper left panel). The values shown are normalized with 18S rRNA level and expressed as copies per μg RNA (n = 3). Electrophoretic gel images of qPCR products (lower left panel) and PCR primer location in the *IP6K2* genomic locus (right panel) was also depicted. W, WT; K, *IP6K2*^-/-^; n.d., not detected; *, non-specific band. **C.** Representative Western blot image of IP6K1 and IP6K2 expression in the cerebrum of *IP6K2*^-/-^ and WT mice. β-actin was used as the internal control. **D-F.** The concentrations of IP_6_ **(D)** and IP_7_ **(E)**, and IP_7_/IP_6_ ratios **(F)** in the CNS, GIT, and other organs of *IP6K2*^-/-^ and WT mice. The values shown represent the mean□±□SD of five independent experiments and are expressed as pmol per mg of organ weight. Asterisks indicate statistical significance (P□<□0.05, Student’s *t*-test) compared with WT mice. **G, H.** Representative SRM chromatograms of IP_6_ (upper panel) and IP_7_ (lower panel) in the stomach **(G)** and duodenum **(H)** of *IP6K2*^-/-^ and WT mice. The three best transitions per molecule are shown for peak identification of each compound. Arrows indicate the SRM peak of each analyte.

### IP6K2-dependent enhanced IP_7_ metabolism exists in the gut and duodenal muscularis externa where the myenteric plexus is located

Since IP_7_-synthesizing kinase IP6K2 is selectively expressed in enteric neurons (Fig. 4), we next sought to investigate IP_7_ metabolism in the mammalian ENS. To this end, we collected the stomach and the consecutive 5 cm segments of duodenum, jejunum, and ileum from standard diet-fed or fasted mice. Some of these organs collected were subsequently used to isolate the muscularis externa where the myenteric plexus is located. These total GIT tissues and their muscularis externa were subjected to HILIC-MS/MS analysis to compare their IP_7_ metabolism (Fig. 6A). Similar to the results shown in Fig. 3, 48 h fasting of mice rendered drastic reduction of IP_6_ and IP_7_ levels with concomitant increase of IP_7_/IP_6_ ratio in total GIT tissues (Fig. 6B-D). Although the muscularis externa contained less IP_6_ and IP_7_ than total GIT tissues, the muscle layer exhibited a higher IP_7_/IP_6_ ratio than total GIT tissues, which was less dependent on dietary conditions. IP_7_/IP_6_ ratio of the duodenal muscularis externa was highest among the corresponding muscle layers of the neighboring GITs, implying highly active IP_7_ metabolism in the duodenal ENS. To verify the relationship between IP6K2-IP_7_ axis and ENS, we first attempted to visualize the duodenal myenteric plexus of *IP6K2*^-/-^ mice by whole mount immunostaining (Fig. 6E). We found that *IP6K2* deletion largely affected neither the morphological features nor the neuronal cell density in the duodenal myenteric plexus (Fig. 6F). We next prepared the muscularis externa from the stomach to the ileum of WT and *IP6K2*^-/-^ mice, first depleting dietary IP_7_ in the GIT by 48 h fasting and performed HILIC-MS/MS analysis to evaluate IP_7_ metabolism in the ENS of *IP6K2*^-/-^ proximal GITs (Fig. 6G). While IP_6_ levels in the muscularis externa were almost equivalent between WT and *IP6K2*^-/-^ mice, IP_7_ levels and IP_7_/IP_6_ ratios were significantly reduced in the gut and duodenal muscularis externa of *IP6K2*^-/-^ mice (Fig. 6H-J). These results suggest that IP6K2 actively produces IP_7_ in the gut and duodenal muscularis externa where enteric neurons are concentrated.

**Figure 6.**
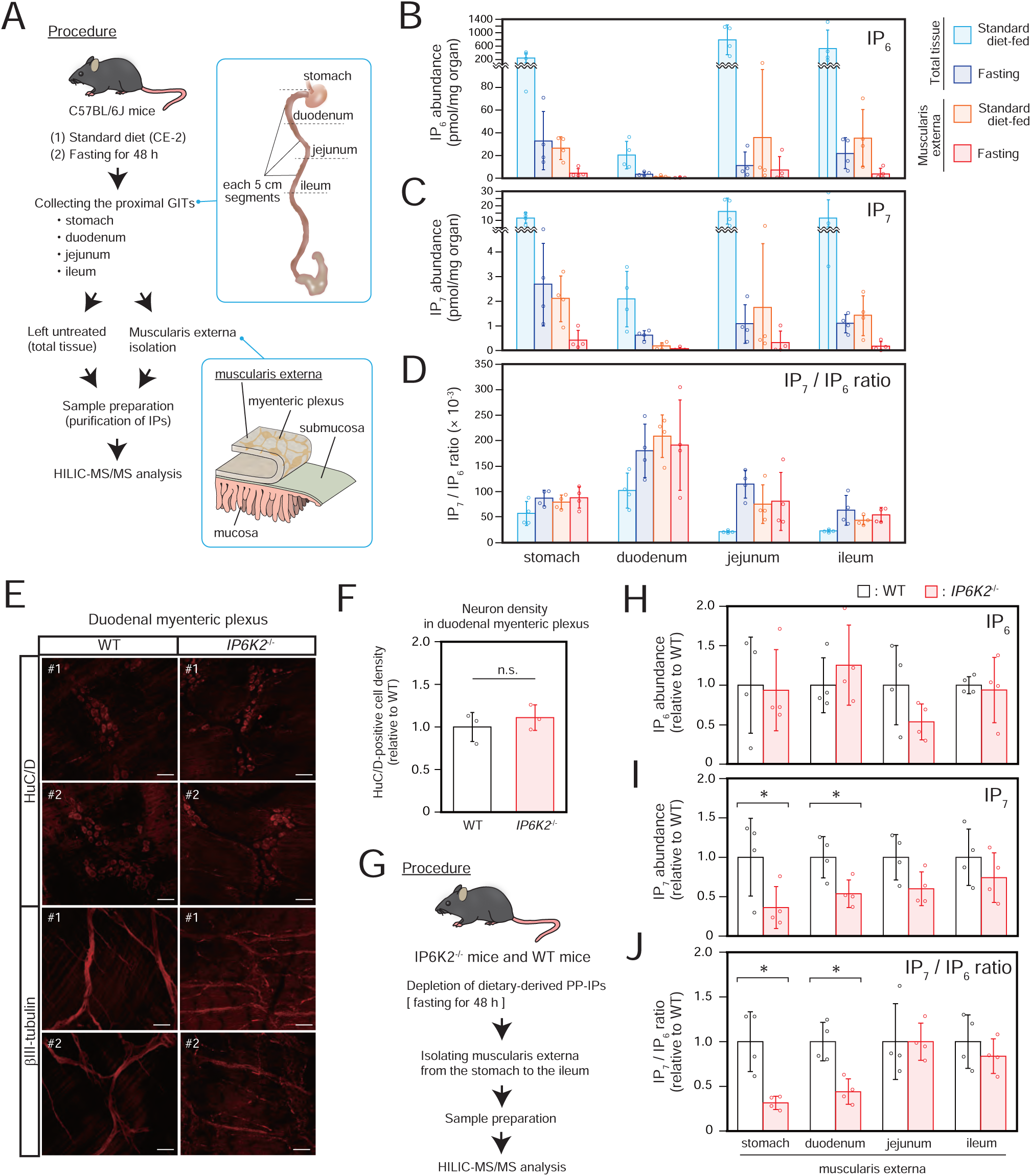
IP6K2-dependent enhanced IP_7_ metabolism exists in the gut and duodenal muscularis externa where the myenteric plexus is located. **A.** Schematic illustration of the experimental workflow. C57BL/6J mice were fed a standard diet, or fasted for 48 h. These mice were sacrificed to collect four stomach and 3 consecutive 5-cm segments of the proximal GIT (duodenum, jejunum, ileum). The muscularis externa containing myenteric plexus as well as total tissues in the proximal GITs were subjected to HILIC-MS/MS analysis. **B-D.** The concentrations of IP_6_ **(B)**, IP_7_ **(C)**, and IP_7_/IP_6_ ratios **(D)** in the muscularis externa and total tissue of four proximal GIT segments of C57BL/6J mice under the two different conditions. The values shown represent the mean□±□SD of four independent experiments and are expressed as pmol per mg of organ weight. **E.** Whole mount immunostaining of WT and *IP6K2*^-/-^ duodenal muscularis externa using anti-neuronal markers antibodies. Two different areas of confocal microscopic images of each neuron marker are shown. The neuronal markers HuC/D and βIII-tubulin were detected to identify enteric neuronal somas and enteric nerve fibers in the myenteric plexuses, respectively. Scale bar = 50 μm. **F.** The concentration of enteric neurons in WT and *IP6K2*^-/-^ duodenal muscularis externa. The values shown represent the mean ±□SD of three independent experiments and are expressed relative to those of WT mice. n.s., not significant (Student’s *t*-test). **G.** Schematic illustration of the experimental workflow. *IP6K2*^-/-^ and WT mice fasted for 48 h were sacrificed to collect four proximal GIT segments (stomach, duodenum, jejunum, ileum), which were then subjected to isolate muscularis externa. **H-J.** The abundances of IP_6_ **(H)** and IP_7_ **(I)**, and IP_7_/IP_6_ ratios **(J)** in the muscularis externa of the four GIT segments of *IP6K2*^-/-^ and WT mice. The values shown represent the mean□±□SD of four independent experiments and are expressed relative to those for WT mice. Asterisks indicate statistical significance (P□<□0.05, Student’s *t*-test) compared with WT mice.

### The IP6K2-IP_7_ axis is crucial for certain neurotranscriptome profiles associated with ENS development and functioning

Considering the active IP6K2-IP_7_ axis in the ENS, we assumed that alteration of IP_7_ metabolism by *IP6K2* deletion might affect neuronal status in the proximal GIT. Thus, we randomly selected two neuronal genes expressed in the GIT as well as the CNS (Gremel et al, 2015), namely *dopamine receptor D5* (*Drd5*), and *cholecystokinin B receptor* (*Cckbr*), and investigated their mRNA levels in both the CNS and GIT by quantitative PCR (qPCR) (Fig. 7A). Compared with those in WT mice, these mRNA levels were explicitly increased from the stomach through the small intestine of *IP6K2*^-/-^ mice, especially in the duodenum, but not the colon and CNS. To comprehensively appreciate the role of IP6K2-dependent IP_7_ metabolism in neuronal gene regulation in the mammalian ENS, we isolated the duodenal muscularis externa from WT and *IP6K2*^-/-^ mice and performed whole transcriptome analysis by RNA-sequencing (RNA-seq) (Fig.7B). Gene set enrichment analysis (GSEA) showed that *IP6K2* deletion suppressed certain gene sets associated with neural stem/progenitor cells, oligodendrocyte progenitor cells, and glial cells, concomitantly with the induction of those of mature neurons such as inhibitory, dopaminergic or GABAergic neurons (Fig. 7C, D and Data S1), implying that inhibition of the IP6K2-IP_7_ pathway triggers neurodevelopmental imbalance in the mammalian ENS. The RNA-seq analysis also exhibited that 107 and 134 out of 23,405 genes were more than 1.5-fold increased or decreased in *IP6K2*^-/-^ with P-value less than 0.05, respectively (Fig. S4A). Pathway enrichment analysis of these genes showed that transcripts increased more than 1.5-fold in *IP6K2*^-/-^ were significantly enriched for proteins involved in neuronal signaling (neuroactive ligand-receptor interaction of KEGG annotation) (Fig. S4B). In these transcripts, we observed that 7 genes associated with neuronal function (*Nckipsd*, and *Hrh4*) or development (*Noto*, *Tbx1*, *Tbx18*, *Pax7*, and *Mycn*) were prominently altered in their transcript levels between WT and *IP6K2*^-/-^ (Fig. 7E). To validate our RNA-seq results, differential expression of these 7 neuronal genes were further assessed by qPCR and all of these candidate genes exhibited similar significant or prominent changes in transcript levels as observed in RNA-seq results (Fig. 7F). Quantitative PCR analysis also showed that expression of other neuronal genes, including *Drd5* and *Cckbr*, explicitly increased in *IP6K2*^-/-^ duodenal muscularis externa (Fig. 7G). These changes were not observed in the RNA-seq analysis possibly because they were below the lower detection limit and/or quantitation error (Robert and Watson, 2015; Everaert et al, 2017). Collectively, the IP6K2-IP_7_ axis contributes to certain neurotranscriptome profiles involved in ENS development and functioning.

**Figure 7.**
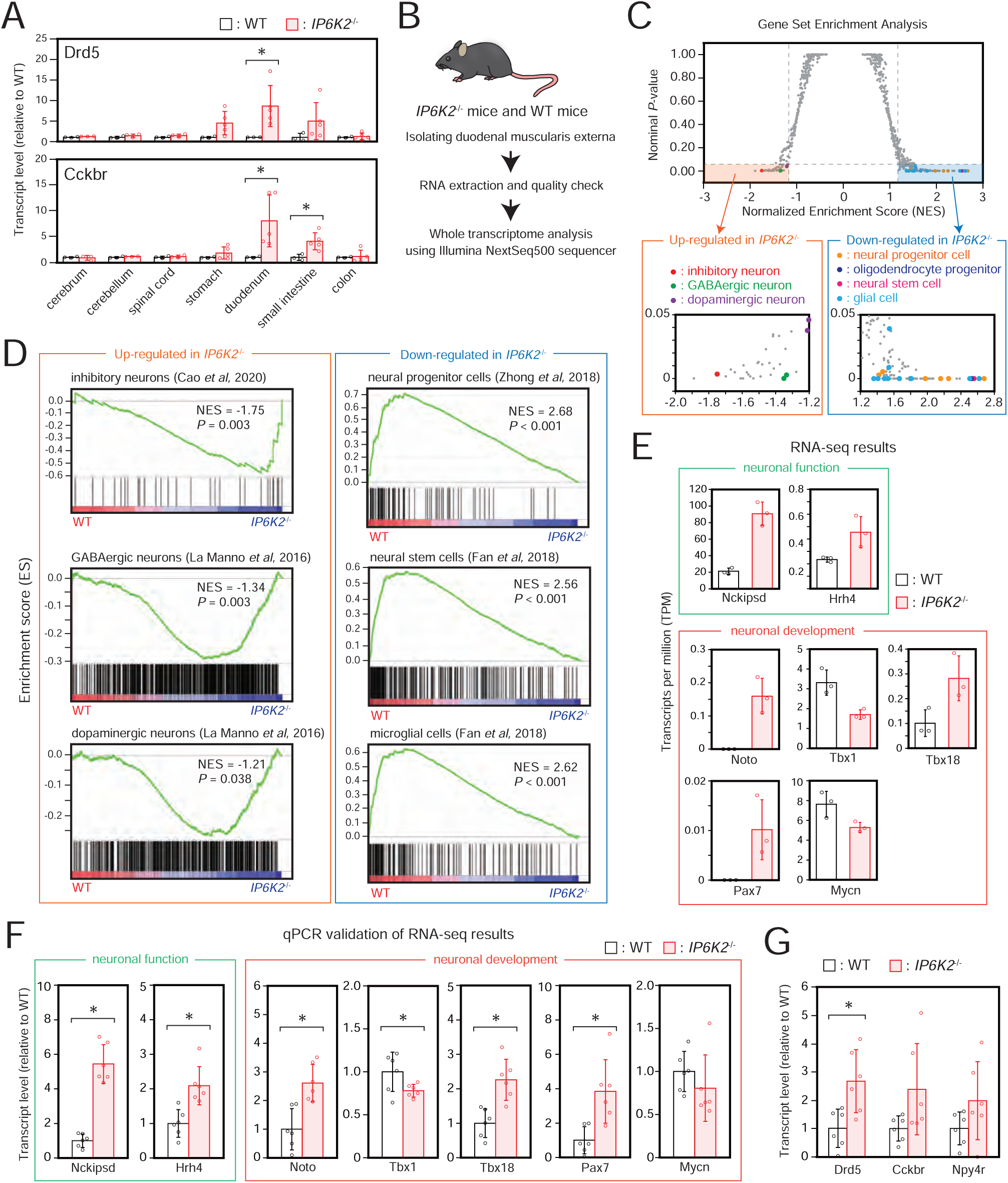
IP6K2-IP_7_ axis is crucial for certain neurotranscriptome profile associating with ENS development and functioning. **A.** Transcript levels of two different neuronal genes (*Ddr5* and *Cckbr*) in the CNS and GIT of *IP6K2*^-/-^ and WT mice. Data were normalized to 18S rRNA level. The values shown represent the mean□±□SD of three (CNS of *IP6K2*^-/-^, and CNS and GIT of WT mice) and five (GIT of *IP6K2*^-/-^ mice) independent experiments and are expressed relative to those of WT mice. Asterisks indicate statistical significance (P□<□0.05, Student’s *t*-test) compared with WT mice. **B.** Schematic illustration of the experimental workflow. *IP6K2*^-/-^ and WT mice were sacrificed to collect the duodenal muscularis externa. High-quality total RNAs isolated from these tissues (each n =3) were subjected to whole transcriptome analysis by high-throughput RNA sequencing. **C.** Gene Set Enrichment Analysis (GSEA) of the enriched gene signature in *IP6K2*^-/-^ duodenal muscularis externa. Cell type signature gene sets (C8 in The Molecular Signatures Database ver7.5.1; http://www.gsea-msigdb.org/gsea/msigdb/index.jsp) were used for this analysis. Horizontal dashed line indicates nominal P-value 0.05, and vertical lines indicate normalized enriched score (NES)□± 1.2 cutoff. Gene sets assigned to neural progenitor cells and oligodendrocyte progenitor cells (source data is derived from Zhong et al, 2018), neural stem cells and glial cells (Fan et al, 2018), and mature neurons (inhibitory neurons, Cao et al, 2020; GABAergic and dopaminergic neurons, La Manno et al, 2016) with nominal P < 0.05 and NES > 1.2 or < -1.2 are labeled in colored dots. **D.** Representative GSEA plots of the gene sets enriched among up-regulated (inhibitory neurons, GABAergic neurons, dopaminergic neurons) or down-regulated (neural progenitor cells, neural stem cells, glial cells) genes by genetic ablation of *IP6K2* in the duodenal muscularis externa. **E.** Normalized expression levels (transcripts per million, TPM) from RNA-seq data for 7 neuronal genes prominently and significantly (P < 0.05, Student’s *t*-test) accumulated or depleted in *IP6K2*^-/-^ duodenal muscularis externa compared with WT counterparts (n = 3). **F.** Validation of RNA-seq results by qPCR. Data were normalized to β-actin level. The values shown represent the mean□±□SD of six independent experiments and are expressed relative to those for WT mice. Asterisks indicate statistical significance (P□<□0.05, Student’s *t*-test) compared with WT mice. **G.** Transcript levels of three different neuronal genes (*Ddr5*, *Cckbr* and *Npy4r*) in the duodenal muscularis externa of *IP6K2*^-/-^ and WT mice. Data were normalized to β-actin level. The values shown represent the mean□±□SD of six independent experiments and are expressed relative to those for WT mice. Asterisks indicate statistical significance (P□<□0.05, Student’s *t*-test) compared with WT mice.

## Discussion

Mammalian PP-IPs have been implicated in obesity and diseases such as cancer and neurodegenerative disorders, and thus, their metabolism is a promising drug target (Shears, 2016; Chakraborty, 2018). For this reason, *in vivo* PP-IP profiling of mammalian tissues is an important subject of research. However, this objective has been thwarted by various technical difficulties. Recently, we developed an HILIC-MS/MS analysis protocol for the sensitive and specific detection of IP_7_ and its precursor IP_6_ (Ito et al, 2018). In this study, we quantified *in vivo* PP-IP levels in mammalian organs using a refined HILIC-MS/MS protocol and evaluated the contribution of IP6K2 to PP-IP metabolism by analyzing mice lacking this IP_7_-synthesizing kinase.

We observed abundant IP_6_ and a small but detectable quantity of IP_7_ in various mammalian organs. Specifically, a discernible level of IP_7_ was detected in the mammalian CNS. These results correlate with the fact that expression of the IP_7_-synthesizing kinases IP6K1 and IP6K2 is ubiquitous and highest in the CNS (Saiardi et al, 2001; Moritoh et al, 2016; Laha et al, 2021). Since IP_7_ levels in the CNS remain constant irrespective of dietary supply (Fig. 3C and H), it is plausible that food-derived IP_7_ is not directly delivered to the CNS. In agreement with this idea, previous studies on rodents have reported that food-derived IPs are degraded in the GIT and released into the circulation as myo-inositol or inositol monophosphate (IP_1_) (Sakamoto et al, 1993; Eiseman et al, 2011). There is also evidence that circulating plasma contains no higher order IPs, such as IP_6_ (Wilson et al, 2015). Considering our observation that IP_7_ levels were significantly decreased in the CNS of *IP6K2*^-/-^ mice compared with those in WT counterparts (Fig. 5E), it is reasonable to regard the IP_7_ detected in the CNS as endogenously generated. Intriguingly, the IP_7_/IP_6_ ratio in the spinal cord was higher than those in the cerebrum and cerebellum, suggesting heterogeneous IP_7_ metabolic activity in the rostral and caudal CNS.

Surprisingly, we found that standard diet-fed rodents had far more IP_7_ in the GIT than in the CNS (Fig. 2B). Furthermore, the IP_7_/IP_6_ ratio, an indicator of PP-IPs metabolism, was far higher in the GIT than in the CNS (Fig. 2C). The mouse diet affected the level of IP_6_ and IP_7_ in the GIT but did not influence the high IP_7_ metabolism, as revealed by the IP_7_/IP_6_ ratio (Fig. 3E and I). The dietary influence on IP_6_ and IP_7_ levels in the GIT is likely a direct consequence of the availability of inositol in plant-derived food (CE-2). Some of this inositol could be generated from IP_6_ by intestinal flora (Priyodip et al, 2017) and directly absorbed by inositol transporters such as SMIT1, SMIT2, and HMIT (Schneider, 2015). A considerable amount of IP_7_ was detected in the GIT of rodents even when the supply of dietary PP-IPs was almost depleted (Fig. 3C and H). The IP_7_/IP_6_ ratio was heterogeneous along the GIT but significantly higher in the proximal GIT of these dietary PP-IP-depleted rodents (Fig. 3E and I), indicating that substantial endogenous IP_7_ metabolism occurs in the proximal GIT. Accordingly, IP_7_ levels in the stomach and duodenum were significantly diminished in *IP6K2*^-/-^ mice under dietary PP-IP-depleted conditions (Fig. 5E). Therefore, our HILIC-MS/MS analysis unexpectedly revealed enhanced IP_7_ metabolism in the mammalian GIT.

The GIT consists of several histological layers including the muscularis externa that contains the myenteric plexus, a collection of large neuronal assemblies in the GIT. Our HILIC-MS/MS survey of the proximal GIT clarified that the muscularis externa has a higher IP_7_/IP_6_ ratio than whole GIT tissues, and the duodenal muscle layer has a much higher IP_7_/IP_6_ ratio than those of neighboring GIT segments (Fig. 6D). Considering the expression of the IP_7_-synthesizing enzyme IP6K2 in the myenteric plexus (Fig. 4A-C) and the significant decrease of IP_7_/IP_6_ ratio in *IP6K2*-deficient gut and duodenal muscle layers (Fig. 6J), these observations lead to the idea that IP6K2 actively synthesizes endogenous IP_7_ in the ENS of the proximal GIT. Our results also implied the presence of endogenous IP_7_ in other GIT layers since total GIT tissues of dietary IP_7_-depleted (fasted) mice contained a greater amount of IP_7_ than their corresponding muscle layers (Fig. 6C). Since another major nerve plexus exists in the submucosal layer (*i.e*. submucosal plexus), the submucosal layer may contain endogenous IP_7_ to some extent. This PP-IP might also exist in mucosal epithelium because certain enteroendocrine cells, including Tuft cells, express IP6Ks at the relatively high level (Fig. S3B and C). Tuft cell is one of rare cell types present in intestinal epithelium. Park et al. recently showed that Tuft cell development is controlled by inositol polyphosphate multikinase (IPMK), an enzyme responsible for driving IP metabolic pathway leading to IP_7_ synthesis (Park et al, 2022). This fact and our data (Fig. S3B and C) encourage to deem that IP6K2 and IP_7_ might underlie Tuft cell physiology as well. Future study is required for assessing cell type-specific IP_7_ metabolism both in enteric neurons as well as in other GIT cells to more precisely characterize IP_7_ metabolism in the GIT.

Although *IP6K2* was initially cloned as a Pi uptake stimulator from a rabbit duodenum complementary DNA (cDNA) library (Norbis et al, 1997) and was annotated soon after as encoding an IP_7_-synthesizing enzyme (Saiardi et al, 1999; Schell et al, 1999), the role of IP6K2 in the GIT has not been investigated until now. In this study, we observed that IP6K2 is prominently expressed in myenteric plexus (Fig. 4A-C), and genetic deletion of *IP6K2* diminishes IP_7_ metabolism in the proximal GIT (Fig. 5F) as well as its muscularis externa containing myenteric plexus (Fig. 6J), suggesting the presence of an active IP6K2-IP_7_ pathway in the ENS of the proximal GIT. Our RNA-seq analysis of the duodenal muscularis externa indicated that genetic ablation of *IP6K2* causes certain gene products associated with mature neurons to accumulate concomitantly with the reduction of those of neural progenitor/stem cells and glial cells (Fig. 7C and D). Given that the developmental lineage of enteric neurons comprises several differentiation points such as neural crest cell migration, neuron-glia bifurcation, and neural stem/progenitor cell differentiation into mature enteric neurons (Rao and Gershon, 2018), inhibition of the IP6K2-IP_7_ axis possibly causes developmental imbalances of the ENS at the several differentiation points at least including the maturation of both enteric neurons and glial cells. This idea is also supported by our findings that IP6K2 inhibition significantly altered the expression levels of several transcription factors regulating neural crest cell differentiation (Fig. 7E and F) (Knoepfler et al, 2002; Vitelli et al, 2002; Abdelkhalek et al, 2004; Bussen et al, 2004; Basch et al, 2006; Simões-Costa et al, 2012). In fact, IP6K2 activity was shown to be required for normal migration and development of neural crest cells in zebrafish (Sarmah and Wente, 2010). Besides, genetic inhibition of *IP6K2* in the duodenal muscularis externa significantly or prominently changed mRNA levels of several genes modulating neuronal functions (Fig. 7E, F and G). Notably, *Nckipsd* transcript, one of the transcripts most significantly induced in *IP6K2*^-/-^ duodenal muscularis externa, contributes to the formation of neural dendrites (Fukuoka et al, 2001; Lee et al, 2006) and intracellular neuronal signaling (Kim et al, 2009). These pieces of knowledge lead to the hypothesis that the IP6K2-IP_7_ axis might directly or indirectly contribute to development and several distinct neuronal functions of enteric neurons, even though the axis does not largely affect the entire morphological output of the ENS (Fig. 6E and F).

Future studies are required for elucidating how these differentially-expressed transcripts controlled by the IP6K2-IP_7_ axis individually affect ENS development and functioning. Developmental and functional ENS defects often result in fatal congenital disorders (Furness, 2012; Wright et al, 2021), but *IP6K2*^-/-^ mice do not show such severe phenotypic defects: *IP6K2*^-/-^ mice are born at Mendelian ratio and grow normally, similar to WT mice (Rao et al, 2014). Thus, the IP6K2-IP_7_ axis might serve as a fine-tuning factor for the developmental and functional regulation of the ENS, although we could not exclude the possibility that *IP6K1*, another major IP6K isoform, compensates for the loss of *IP6K2*. It will be meaningful to see whether ENS-specific inhibition of IP6K2 and/or IP6K1 influences gastrointestinal pathophysiologies and development of CNS diseases. Taken together, our observations provide valuable insights into the field of PP-IP biology and neurogastroenterology.

Since dysregulation of IP_7_ metabolism links to various human diseases including neurodegenerative diseases, studying IP_7_ metabolism in human organs provides essential knowledge from the clinical point of view. The refined HILIC-MS/MS protocol we described in this study is capable of detecting IP_6_ and IP_7_ not only in rodent organs but also in human postmortem organs dissected after forensic intervention (Fig. S5). Unlike in rodents, IP_7_ level and IP_7_/IP_6_ ratio in the human proximal GITs (esophagus, greater curvature and lesser curvature of the stomach) were less abundant compared with those in human CNS. This is probably due to the high turnover rate of PP-IPs and the delay in dissecting human postmortem organs. Forensic intervention and subsequent organ dissection take hours after death The presence of the intestinal flora may also facilitate the decomposition of these molecules in the GITs (Musshoff et al, 2011). Thus, care should be taken to assess IP_7_ metabolism in human GITs. Although the refined HILIC-MS/MS protocol can detect both IP_7_ and IP_8_, this protocol failed to detect endogenous IP_8_ in all rodent and human organs examined in this study, even in mouse GIT where IP_7_ was explicitly abundant. This fact suggested that mammalian-derived IP_8_ is far less abundant than IP_7_, and its quantitative evaluation requires sample pooling or a more sensitive analytical protocol such as CE-MS (Qiu et al, 2020). In any case, we demonstrated that our novel protocol was able to evaluate IP_7_ metabolism in human organs. Therefore, we foresee the diagnostic potential of our new analytical technique for analyzing IP_6_ and IP_7_ levels in clinical biopsy.

In conclusion, we investigated the distribution of PP-IPs in mammalian organs using a refined HILIC-MS/MS protocol and demonstrated that IP6K2-dependent IP_7_ metabolism was enhanced in the ENS of the proximal GIT. This finding was corroborated by the observation that impairment of IP6K2-dependent IP_7_ metabolism significantly altered certain neurotranscriptome profiles involved in ENS development and functioning. Further studies are needed to dissect molecular mechanisms underlying IP6K2-IP_7_ axis-mediated neurotranscriptional regulation in the ENS, the role of IP_7_ in neurogastroenterology, and processes involving the gut-brain axis. We believe that these findings shed new light on the physiological significance of the mammalian PP-IP pathway as well as the regulatory mechanisms of ENS functioning, which might contribute to a better understanding of human diseases associated with altered PP-IP metabolism.

## Acknowledgements

We greatly appreciate Prof. Solomon H. Snyder of Johns Hopkins University for providing *IP6K2*^-/-^ mice, Prof. Kazunori Nakajima and Dr. Yuki Hirota of Keio University for fruitful discussion, and Prof. Shinji Hadano of Tokai University for providing organs of purified diet-fed Sprague–Dawley rats. We thank Mr. Shingo Utsuki of the Liaison Laboratory Research Promotion Center in Kumamoto University for technical support of RNA sequencing, and the following staff members of the Support Center for Medical Research and Education of Tokai University for technical assistance: Akemi Kamijo, Katsuko Naito, Sachie Tanaka, Sanae Ogiwara, and Kayoko Iwao for animal experiments; Chisa Okada and Yuka Kitamura for confocal microscopic observation; and Keiko Yokoyama and Sanae Isaki for preliminary microarray experiments and data analysis. We also thank Editage (www.editage.jp) for English language editing. This research was partly supported by a Medical Research Council grant MR/T028904/1 (to A.S.) and grants-in-aid numbers JP19K07851 and JP22K07379 (to E.N.) and JP20H01002 (to M.I.) for scientific research from Japan Society for the Promotion of Science.

## Author contributions

Conceptualization, M.I.; Methodology, M.I.; Investigation, M.I., N.F., S.K., S.H., T.I., K.H.; Formal Analysis, M.T., K.K., D.K.; Resources, C.W., Y.K., H.J.J.; Writing – Original Draft, M.I.; Writing – Review & Editing, M.I., A.S., E.N.; Visualization, M.I.; Funding Acquisition, M.I., A.S., E.N.; Supervision, A.S., E.N.; Project Administration, E.N.

## Competing interests

The authors declare no competing interests.

## STAR Methods

### Key resources table

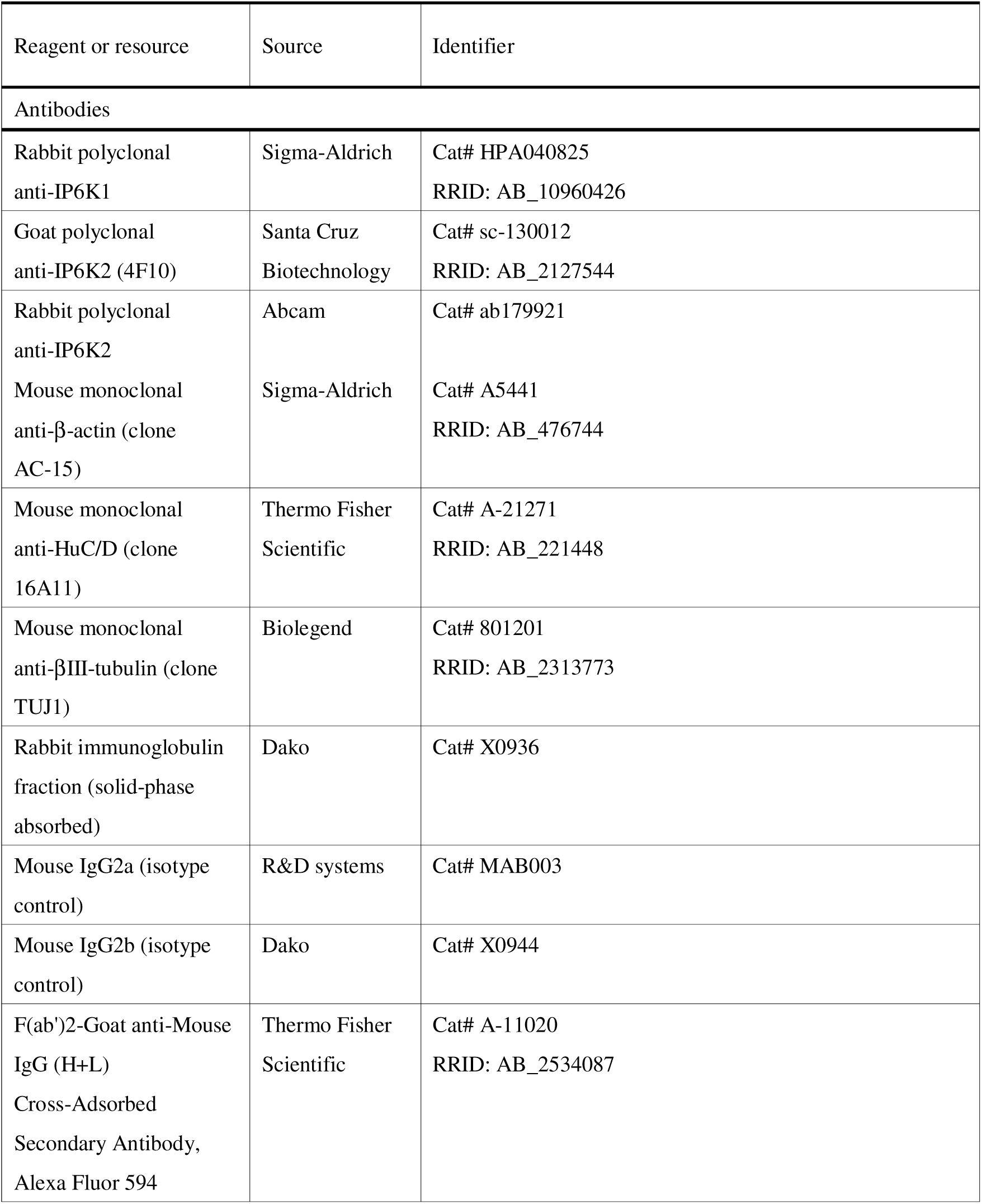
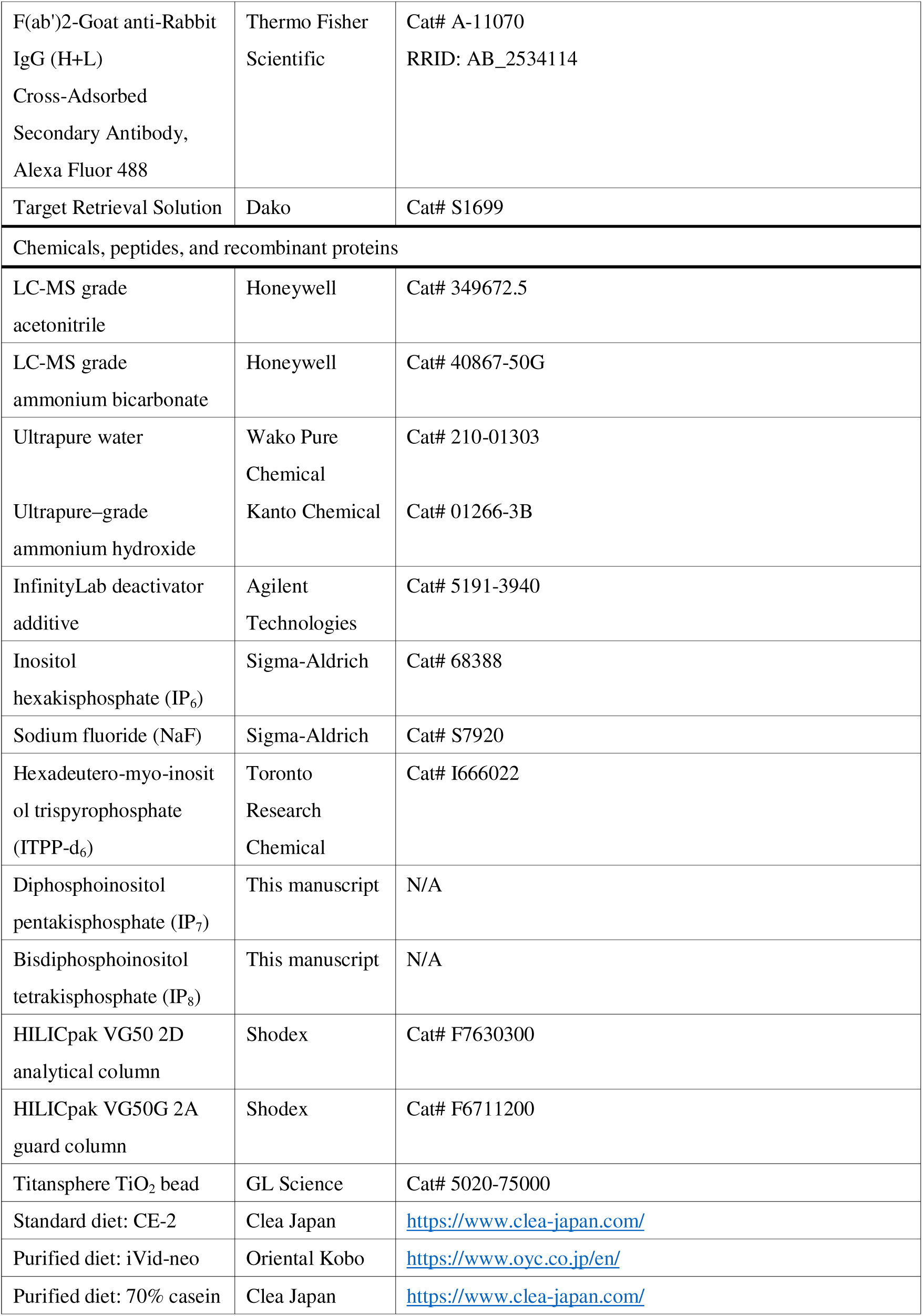
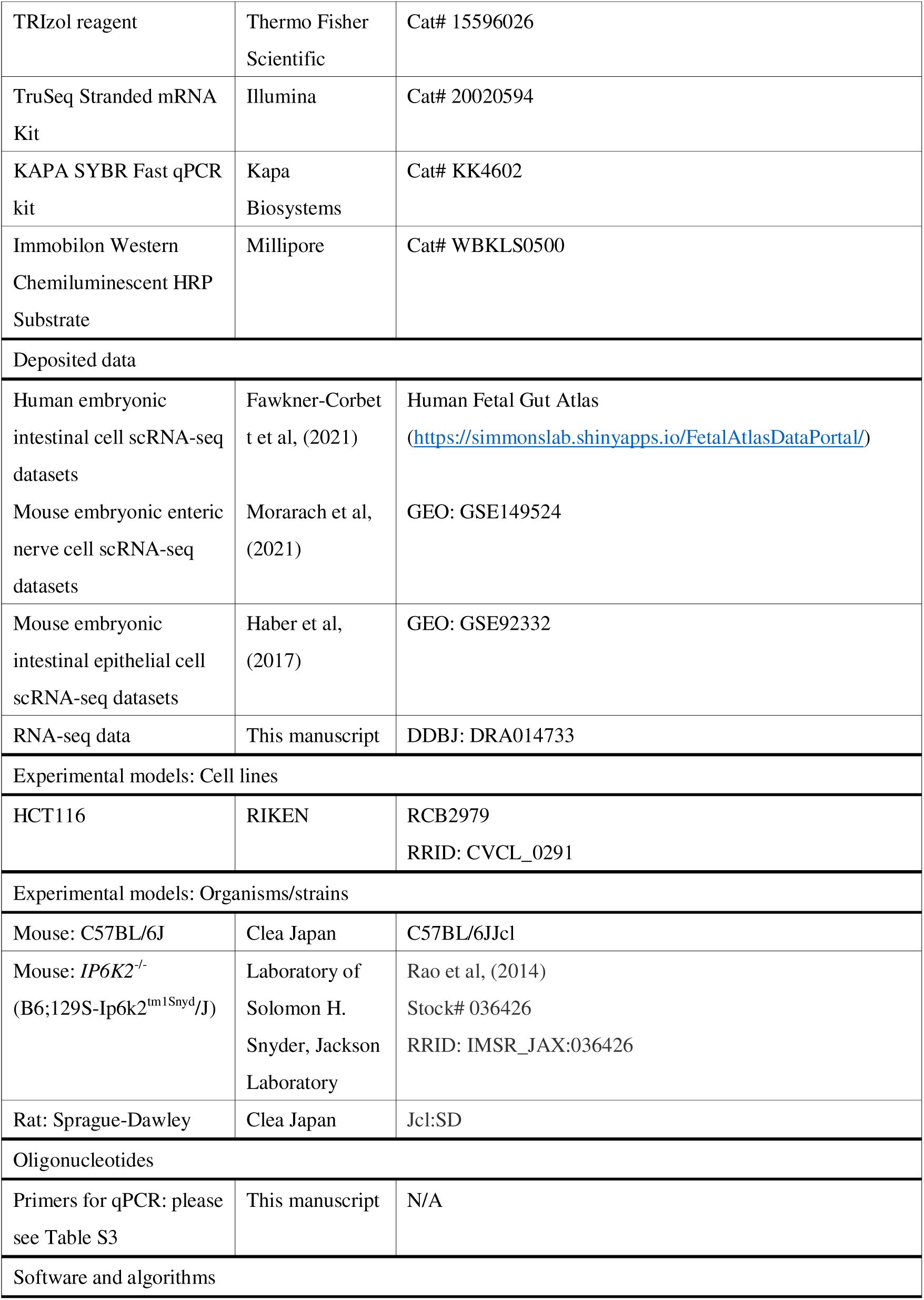
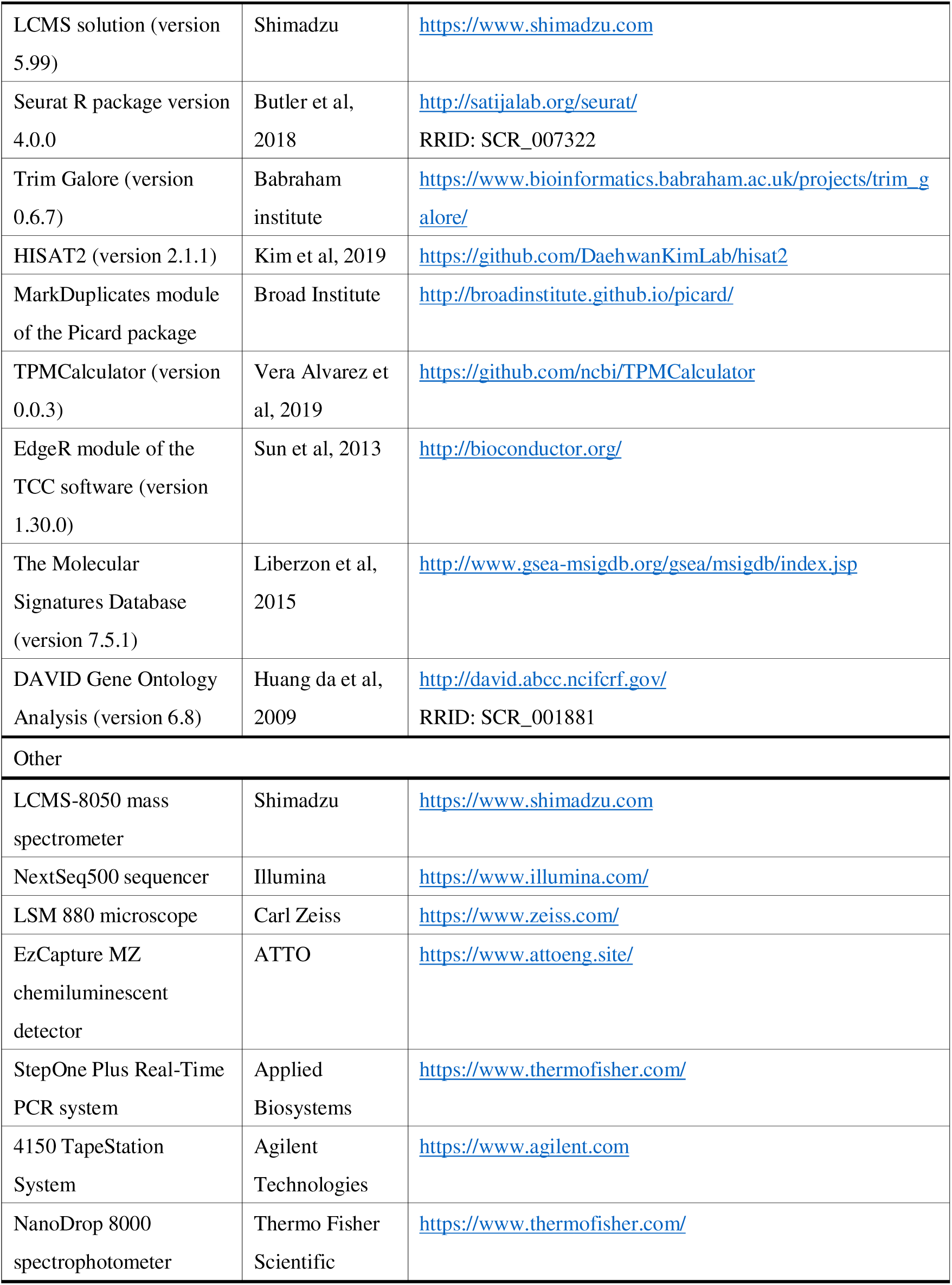

## RESOURCE AVAILABILITY

### Lead contact

Further information and requests for resources and reagents should be directed to and will be fulfilled by the Lead Contact, Eiichiro Nagata or Masatoshi Ito.

### Materials availability

Requests for materials generated in this study should be directed to the Lead Contact. The availability of these materials may be limited because their chemical synthesis requires multiple laborious and costly processes.

### Data and code availability

The raw RNA-seq data have been deposited at the DNA Data Bank of Japan (DDBJ) and are publicly available as of the date of publication (accession number DRA014733). Raw GSEA data shown in Fig. 7C are available in the Data S1 file. Other individual datasets and corresponding files generated in this study are available upon reasonable request from the Lead Contact.

## EXPERIMENTAL MODEL AND SUBJECT DETAILS

### Cell culture

HCT116 cells were cultured in DMEM (Nacalai Tesque, Kyoto, Japan) supplemented with 10% FBS in 5% CO_2_. Cells prepared at 60% confluence in 10 cm dishes were incubated for 1 h with or without 50 mM sodium fluoride (Sigma-Aldrich, St. Louis, MO, USA). After washing twice with PBS, the cells were lysed in cell lysis buffer (0.01% Triton X-100, 1 mM EDTA, 20 mM Tris-HCl). A small aliquot was set aside for protein quantitation, and the rest was used for purification of IPs.

### Mouse organs

All experiments involving animals were performed in accordance with protocols approved by institutional animal care guidelines (Tokai University School of Medicine). Male C57 BL/6J mice and Sprague–Dawley rats obtained from Clea Japan (Tokyo, Japan) were maintained on a standard diet (CE-2; Clea Japan) or purified diet (iVid-neo; Oriental Kobo, 70% casein; Clea Japan). Some mice fed a standard diet or purified diet were fasted for 48 h before sacrifice. *IP6K2*^-/-^ and WT mice maintained on a standard diet were switched to the purified diet for a week and subsequently fasted for 48 h before sacrifice. During fasting, mouse cages were changed to clean ones with new bedding every 24 h to reduce coprophagy. Mice and rats were anesthetized using isoflurane and then sacrificed by whole blood withdrawal from the left atrium. Before dissection of the organs, the animals were perfused transcardially with ice-cold PBS to wash out the residual blood and prevent the detection of IPs derived from blood cells. GIT organs (stomach, duodenum, small intestine, and colon) were cut open to remove feces and then extensively rinsed with PBS to wash out any dietary residuals. The duodenum, small intestine, and colon were harvested by cutting a 5 cm (mouse) or 10 cm (rat) segment from the distal end of stomach, between 10 and 15 cm (mouse) or 20 and 30 cm (rat) away from the duodenum and from the anus, respectively. The harvested organs were frozen until further use.

### Isolation of muscularis externa from mouse GITs

The muscularis externa containing myenteric plexuses was prepared from mouse GITs as previously described with some modification (Fujita et al, 2018; Ahrends et al, 2022). For HILIC-MS/MS, mouse GIT segments were cut open along the attachment line of the mesentery and then placed onto a cold surface with the muscularis externa facing up. The muscularis externa of the GIT segments was isolated by gently scraping the outer layer with watchmaker tweezers under a binocular stereomicroscope. For whole-mount immunostaining, the mouse duodenum was cut open along the mesentery line, pinned onto a rubber plate, and then fixed with 4% paraformaldehyde (PFA) overnight at 4 °C. The muscularis layer was then gently separated from the GIT segment using watchmaker tweezers and a cotton swab under a binocular stereomicroscope. For RNA extraction, mouse GITs were immersed in saturated ammonium sulfate solution containing 20 mM EDTA and 25 mM sodium citrate (pH5.2) to inhibit RNA degradation. The muscularis externa of the segments was placed over a glass rod and then peeled away using a cotton swab along the attachment line of the mesentery under a binocular stereomicroscope as described previously (Smith et al, 2013). Isolated muscularis externa was stored in saturated ammonium sulfate solution and then frozen until further use.

### Human postmortem organs

The human study was approved by the Ethics Committee of Tokai University (institutional review board number: 20I-02), and the study protocol conformed to the ethical guidelines of the 1975 Declaration of Helsinki (World Medical Association, 2013). Written informed consent, allowing the experimental use of the organ samples, was obtained from the relatives of all subjects. Human postmortem organs were obtained at autopsies from three donated bodies (two men and one woman; mean age, 62.3 ± 22.8 years; average body mass index, 26.4 ± 6.6). Some anomalies, such as cardiac hypertrophy, were observed in their bodies by a forensic pathologist. To minimize organ decomposition, organ sampling was confined to cases where the death date and ambient temperature were explicit and the accumulated degree–days [ADD; environmental temperature (°C) × postmortem interval (day)] value—an index for evaluating the quality of forensic samples (Pittner *et al*, 2016)—of all three bodies were very low (close to or less than 20). The harvested organ samples (approximately 400 mg) were frozen until further use.

## METHOD DETAILS

### Gel electrophoresis of synthetic PP-IPs

IP_7_ and IP_8_ were synthesized from myo-inositol using fluorenylmethyl phosphoramidite chemistry as described previously (Pavlovic et al, 2016). The synthetic PP-IPs were validated using polyacrylamide gel electrophoresis as previously described (Losito et al, 2009). Briefly, synthetic PP-IPs samples mixed with orange G and bromophenol blue loading buffer were applied onto 35% polyacrylamide/Tris-borate-EDTA gel. The samples were electrophoresed overnight at 4 °C at 600 V and 6 mA until the orange G and bromophenol blue had run through two-thirds of the gel. Gels were stained with toluidine blue and scanned using a computer scanner.

### Purification of IPs

IPs in biological samples were purified as described previously (Wilson et al, 2015), with some modification. Frozen organs, diets, and feces samples were homogenized using a Shake Master Neo (Bio Medical Science, Tokyo, Japan) in 500 μL of ultrapure water. Feces samples were air-dried overnight before homogenization for accurate comparison of IP_6_ and IP_7_ concentrations with those in the diet. Crude lysate was mixed with an equal volume of 2 M perchloric acid (PCA), incubated on ice for 30 min, and centrifuged to remove tissue debris. After spiking with 3 nmol of ITPP-d_6_ as an internal control, 5 mg of TiO_2_ beads (GL Sciences, Tokyo, Japan) were added to each sample. The beads were incubated at 4 °C for 30 min and washed twice with 1 M PCA, and then 200 μL of 10% ammonium hydroxide was added for IP elution. The elution step was repeated to maximize recovery. The total eluate was dried using a SpeedVac concentrator (Thermo Fisher Scientific, Waltham, MA, USA) and reconstituted in 125 μL of 100 mM ammonium carbonate/40% acetonitrile buffer, 50 μL of which was used for LC-MS.

### HILIC-MS/MS analysis for PP-IPs

Chromatographic experiments were performed using a Nexera UHPLC instrument (Shimadzu, Kyoto, Japan). HILIC-based chromatographic separation of IP_6_, IP_7_, IP_8_, and the internal control ITPP-d_6_ was achieved using a modified version of a previously described procedure (Ito et al, 2018). The mobile phase was composed of 300 mM ammonium bicarbonate buffer (pH 10.5) containing 0.1% InfinityLab deactivator additive (Agilent Technologies) as the aqueous mobile phase (eluent A), and 90% acetonitrile containing 10 mM ammonium bicarbonate buffer (pH 10.5) and 0.1% InfinityLab deactivator additive as the organic mobile phase (eluent B). Eluent B included more than 10% aqueous solvent to prevent polymeric aggregation of the major constituent (medronic acid) in the additive. In the entire LC system, chromatographic stainless steel tube was treated with 0.5% phosphoric acid in 90% acetonitrile overnight before analysis to block undesirable adsorption of analytes on the surface of the inner wall of the tube, while paying attention not to run the solvent into the mass spectrometer. The total flow rate of the mobile phase was 0.4 mL/min. Linear gradient separation was achieved as follows: 0–2 min, 75% B; 2–12 min, 75%–2% B; 12–15 min, 2% B.

### RNA extraction and quantitative PCR analysis

GIT segments and their muscularis externa were carefully collected and subjected to RNA extraction, as previously described (Augereau et al, 2016). Total RNA was extracted using TRIzol reagent (Invitrogen, Carlsbad, CA, USA). RNA concentration and quality were determined using a NanoDrop 8000 spectrophotometer (Thermo Fisher Scientific) and the 4150 TapeStation system (Agilent Technologies), respectively. Complementary DNA was generated using the High-Capacity Reverse Transcription Kit (Applied Biosystems). qPCR was performed using the KAPA SYBR Fast qPCR kit (Kapa Biosystems, Wilmington, MA, USA) and a StepOne Plus Real-Time PCR system (Applied Biosystems, Foster City, CA, USA). The primer sequences used in this study are listed in Table S3.

### RNA sequencing

Total RNA samples of WT and *IP6K2*-deficient duodenal muscle externa with around 7.0 of RNA integrity number were subjected to RNA-seq analysis. RNA sequencing libraries were prepared using TruSeq Stranded mRNA Kit (Illumina, San Diego, CA, USA) according to the manufacturer’s instructions. Each library was sequenced in 1 × 75 bp of single read mode using a NextSeq 500 platform (Illumina). Adapter sequences are removed from sequencing reads using Trim Galore (version 0.6.7; https://www.bioinformatics.babraham.ac.uk/projects/trim_galore/). Sequence reads were aligned to mouse genome (mm10) by HISAT2 (version 2.1.1) (Kim et al, 2018). Duplicate reads were removed using the MarkDuplicates module of the Picard package (version 2.27.3; http://broadinstitute.github.io/picard/). The following genes were excluded before processing for the expression data analysis: highly expressing mucosal digestive enzyme genes (*Amy2a1*, *Amy2a2*, *Amy2a3*, *Amy2a4*, *Amy2a5*, *Amy2b*, *Amy1*, *Try4*, *Try5*, *Try10*) contaminated during the muscularis isolation, mitochondrial genes, and long non-coding RNAs. Expression levels of genes annotated in GENCODE (version M25) were quantitated by TPMCalculator (version 0.0.3) (Vera Alvarez et al, 2019). The software described above was run with the default parameters. Differentially expressed genes were identified by the EdgeR module of the TCC software (version 1.30.0) (Sun et al, 2013). GSEA was performed as described previously (Subramanian et al, 2005). Gene sets used in this study were retrieved from The Molecular Signatures Database (version 7.5.1; http://www.gsea-msigdb.org/gsea/msigdb/index.jsp) (Liberzon et al, 2015). Pathway enrichment analysis was performed using the online database DAVID (http://david.abcc.ncifcrf.gov) (Huang da et al, 2009).

### Western blot analysis

Western blot analysis was performed as previously described (Nagata et al, 2011). Membranes were incubated with anti-IP6K1 (Sigma-Aldrich), anti-IP6K2 (Santa Cruz Biotechnology, Dallas, TX, USA) and anti-β-actin (Sigma-Aldrich) primary antibodies overnight at 4□°C. After rinsing 3 times in PBS containing 0.05% Tween-20, the membranes were incubated with the appropriate secondary antibodies conjugated with horseradish peroxidase (HRP) (donkey anti-rabbit IgG, HRP-linked F(ab’)_2_ fragment or sheep anti-mouse IgG, HRP-linked F(ab’)_2_ fragment; GE Healthcare, Buckinghamshire, UK). The immunoreactivities of the primary antibodies were visualized with Immobilon Western Chemiluminescent HRP Substrate (Millipore, Billerica, MA, USA) and recorded using an Ez-Capture Analyzer (ATTO, Tokyo, Japan).

### Immunohistochemistry

Preparation of formalin-fixed, paraffin-embedded (FFPE) sections was described previously (Nagata et al, 2016). After deparaffinization and rehydration, the mouse tissue sections were incubated with Target Retrieval Solution (Dako, Glostrup, Denmark) at 98 °C for 10 min. Thereafter, the sections were washed thrice with 0.05% Tween-20 in Tris-buffered saline (TBS), blocked using 5% normal goat serum for 15 min, and then incubated with primary antibodies against IP6K2 (1:100, ab179921, Abcam, Cambridge, MA, USA) or HuC/D (1:100, A-21271, Thermo Fisher Scientific) overnight at 4□°C. Rabbit immunoglobulin (Dako) and mouse IgG2b isotype control (Dako) were used to evaluate non-specific binding. After rinsing thrice with 0.05% Tween-20 in TBS, the sections were incubated with secondary goat anti-rabbit IgG Alexa 488 (1:350, A-11070, Thermo Fisher Scientific) and goat anti-mouse IgG Alexa 594 (1:350, A-11020, Thermo Fisher Scientific) antibodies for 30 min at room temperature. The sections were then washed thrice with 0.05% Tween-20 in TBS and mounted using anti-fading medium (12.5 mg/mL DABCO, 90% glycerol, pH 8.8 in PBS). Confocal fluorescence images were obtained using a LSM 880 microscope (Carl Zeiss, Jena, Germany).

### Whole-mount immunostaining

Immunostaining of duodenal muscularis externa was performed as previously described with minor modifications (Fujita et al, 2018; Ahrends et al, 2022). Briefly, duodenal muscularis externa isolated from WT and *IP6K2*^-/-^ mice were blocked with 3% BSA blocking solution containing the corresponding isotype control antibodies for 2 days after fixation with 4% PFA overnight. The muscle layers were then washed with PBS containing 0.05% Triton X-100 and incubated with diluted primary antibodies against HuC/D (Thermo Fisher Scientific) or βIII-Tubulin (Biolegend, San Diego, CA, USA) for 3 days. After rinsing thrice with 0.05% Triton X-100 in PBS, the muscularis externa were then incubated with secondary goat anti-mouse IgG Alexa 594 (1:350, A-11020, Thermo Fisher Scientific) for 3 h at room temperature. The samples were then washed thrice with 0.05% Triton X-100 in PBS and mounted using anti-fading medium (12.5 mg/mL DABCO, 90% glycerol, pH 8.8 in PBS). Fluorescence images were obtained using a LSM 880 confocal microscope (Carl Zeiss).

### Computational analysis of scRNA-seq datasets

Publicly available human embryonic intestine scRNA-seq processed data (Fawkner-Corbett et al, 2021) and mouse embryonic ENS matrix data (Morarach et al, 2021) were downloaded from the Human Fetal Gut Atlas (https://simmonslab.shinyapps.io/FetalAtlasDataPortal/) and the GEO database (identifier: GSE149524), respectively. Mouse embryonic intestinal epithelial cell data (Haber et al, 2017) were obtained from the GEO database (identifier: GSE92332). The above datasets were analyzed using the R package Seurat version 4.0.0 (Butler et al, 2018) to perform dimensionality reduction by uniform manifold approximation and projection and/or generate dot plots showing the relative expression of IP6Ks across different clusters.

## QUANTIFICATION AND STATISTICAL ANALYSIS

Data are expressed as the mean□±□SD. Differences between two or more groups were analyzed using two-tailed Student’s *t*-test or one-way analysis of variance (ANOVA) followed by Bonferroni-type post-hoc test, respectively. In RNA-seq analyses, P values were determined using the corresponding analytical tools. Statistical significance was set at P < 0.05.

## Notes

### Competing Interest Statement

The authors have declared no competing interest.

