## Supplemental Information for "Inositol pyrophosphate profiling reveals regulatory roles of IP6K2-dependent enhanced IP_7_ metabolism in enteric nervous system"

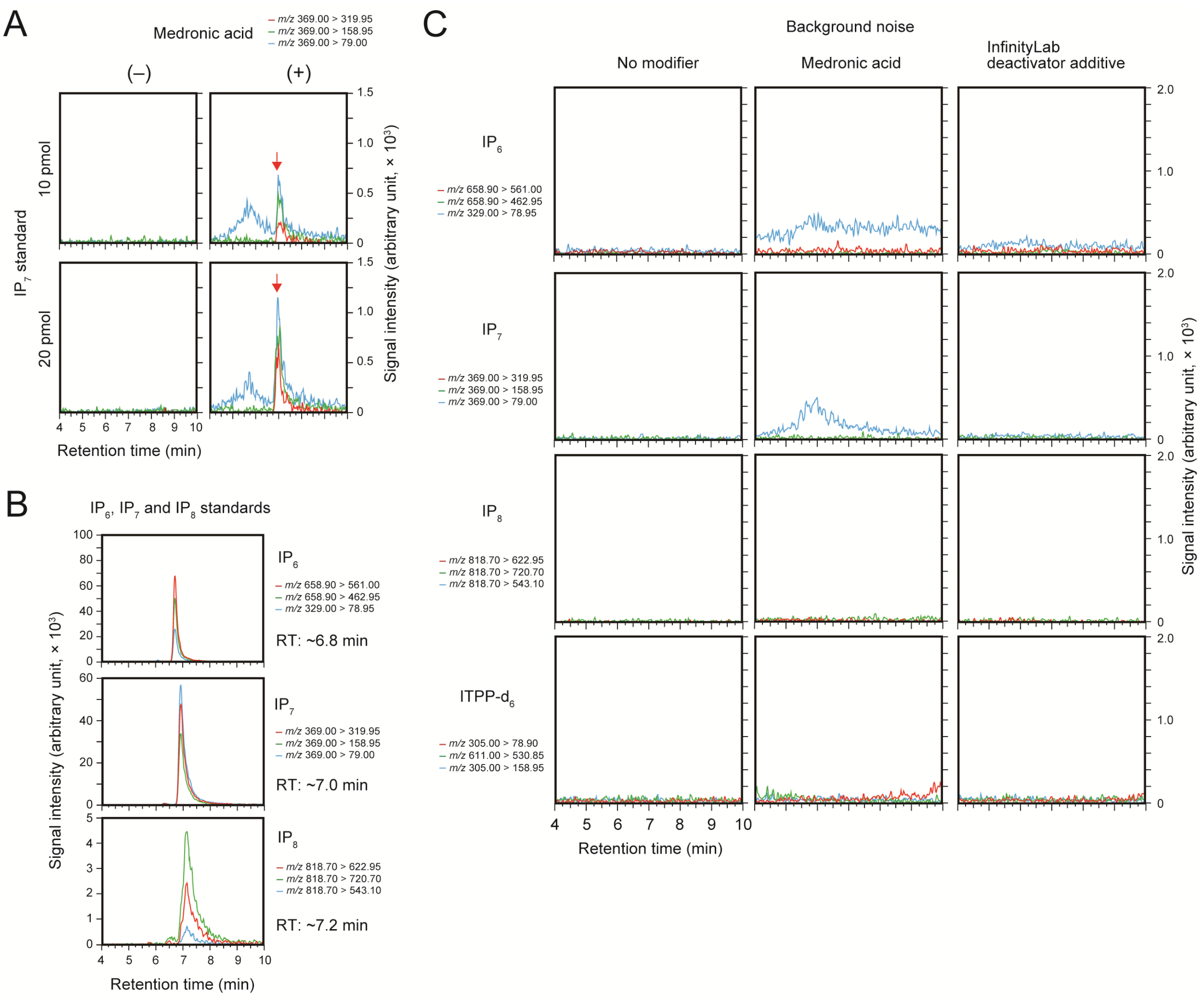


**Figure S1. Effect of medronic acid as a mobile phase modifier on PP-IP analysis (related to Figure 1)**

A. Effect of medronic acid as a mobile phase modifier on the detection of low amounts of synthetic PP-IPs. 10 and 20 pmol of $IP_7$ standard were injected.

B. SRM chromatograms of synthetic $IP_6$, $IP_7$, and $IP_8$ standards. 500 pmol of each standard was injected.

C. Background noise in SRM chromatograms of respective analytes when three different mobile phases were used for HILIC-MS/MS analysis. In SRM transition of $IP_7$, background signals were negligible in the InfinityLab deactivator additive as well as in the absence of a modifier. On the other hand, the background noise was relatively high around the $IP_7$ retention time (~7.0 min) in medronic acid.

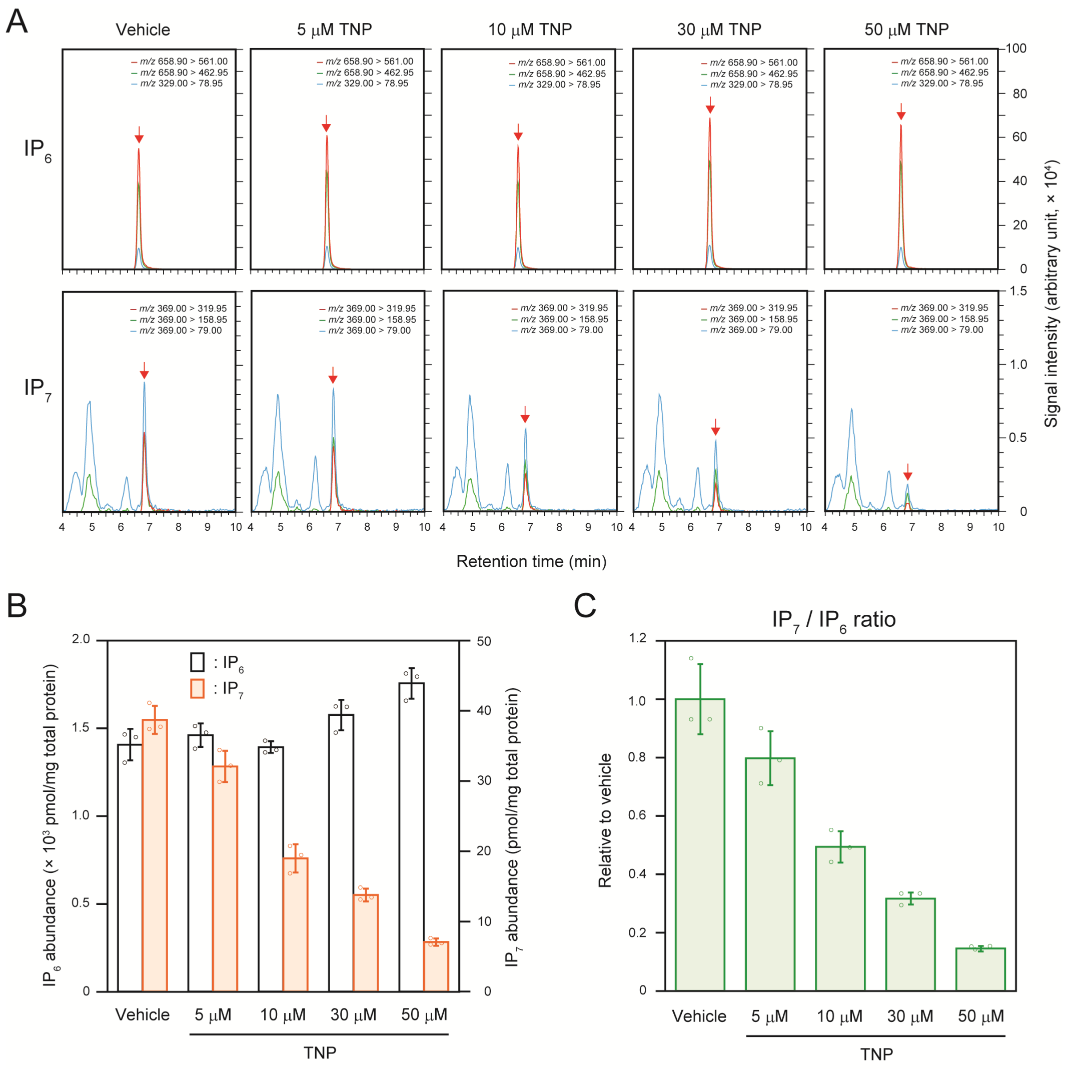


**Figure S2. Inhibition of $IP_7$ production by IP6K inhibitor TNP in HCT116 cells (related to Figure 1)**

A. Representative SRM chromatograms of $IP_6$ and $IP_7$ in HCT116 cells treated for 1 h with different concentrations (0, 5, 10, 30, 50 μM) of the IP6K inhibitor TNP. Arrows indicate SRM peaks of corresponding analytes.

B. Concentrations of $IP_6$ and $IP_7$ in TNP-treated HCT116 cells. The values shown are expressed as pmol per mg of total protein (n = 3).

C. $IP_7/IP_6$ ratios in TNP-treated HCT116 cells. The values shown are expressed relative to those for vehicle-treated counterparts (n = 3).

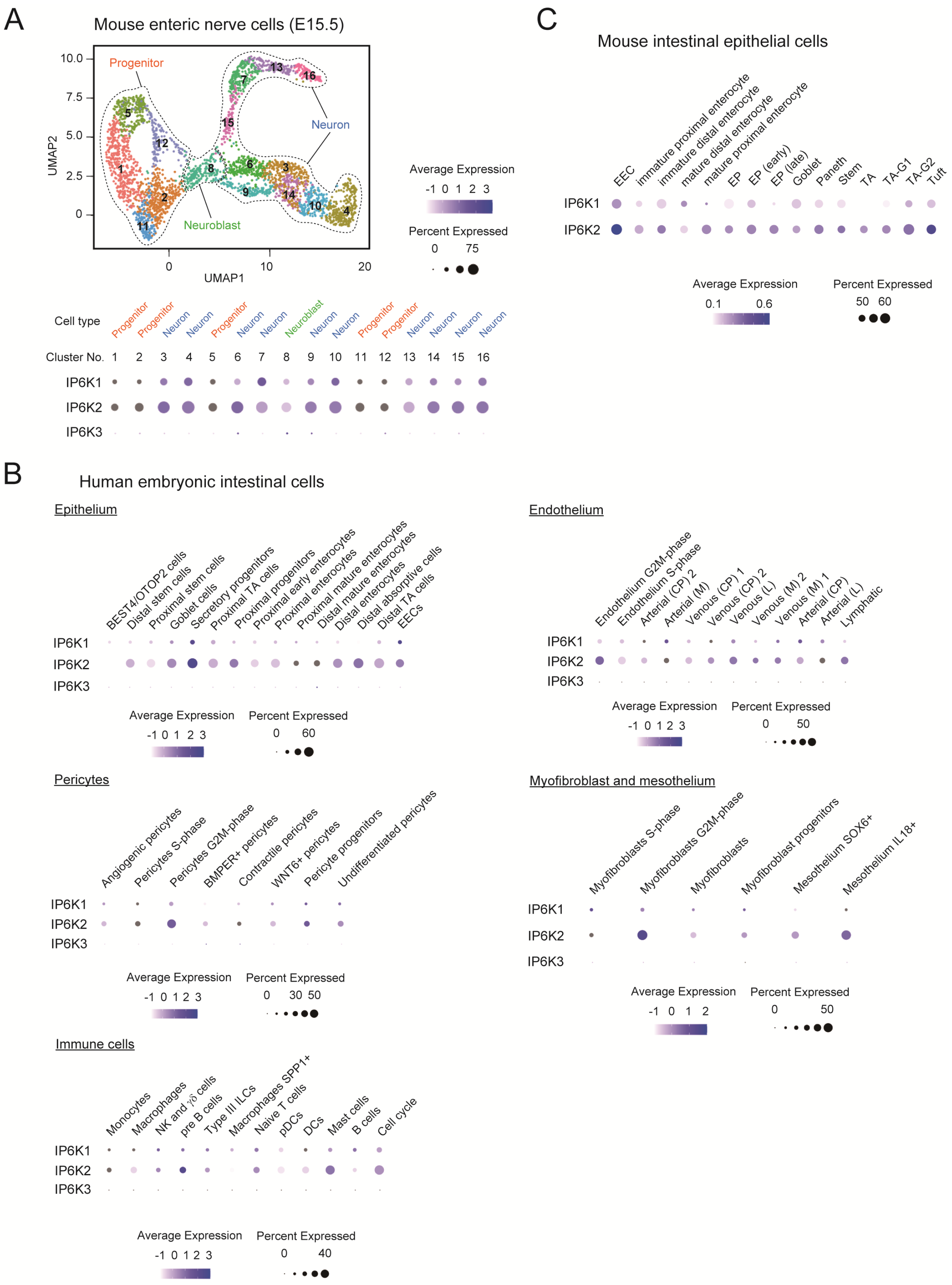
A
Mouse enteric nerve cells (E15.5)
Progenitor
Neuron
Neuroblast
UMAP1
UMAP2
Average Expression
-1 0 1 2 3
Percent Expressed
0 75
Cell type
Progenitor Progenitor Neuron Neuron Progenitor Neuron Neuron Neuroblast Neuron Neuron Progenitor Progenitor Neuron Neuron Neuron Neuron
Cluster No. 1 2 3 4 5 6 7 8 9 10 11 12 13 14 15 16
IP6K1
IP6K2
IP6K3
C
Mouse intestinal epithelial cells
EEC
immature proximal enterocyte
immature distal enterocyte
mature distal enterocyte
mature proximal enterocyte
EP
EP (early)
EP (late)
Goblet
Paneth
Stem
TA
TA-G1
TA-G2
Tuft
IP6K1
IP6K2
Average Expression
0.1 0.6
Percent Expressed
50 60
B
Human embryonic intestinal cells
Epithelium
BEST4/OTOP2 cells
Distal stem cells
Proximal stem cells
Goblet cells
Secretory progenitors
Proximal TA cells
Proximal progenitors
Proximal early enterocytes
Proximal enterocytes
Proximal mature enterocytes
Distal mature enterocytes
Distal enterocytes
Distal absorptive cells
Distal TA cells
EECs
IP6K1
IP6K2
IP6K3
Average Expression
-1 0 1 2 3
Percent Expressed
0 60
Endothelium
Endothelium G2M-phase
Endothelium S-phase
Arterial (CP) 2
Arterial (M)
Venous (CP) 1
Venous (CP) 2
Venous (L)
Venous (M) 2
Venous (M) 1
Arterial (CP)
Arterial (L)
Lymphatic
IP6K1
IP6K2
IP6K3
Average Expression
-1 0 1 2 3
Percent Expressed
0 50
Pericytes
Angiogenic pericytes
Pericytes S-phase
Pericytes G2M-phase
BMPER+ pericytes
Contractile pericytes
WNT6+ pericytes
Pericyte progenitors
Undifferentiated pericytes
IP6K1
IP6K2
IP6K3
Average Expression
-1 0 1 2 3
Percent Expressed
0 30 50
Myofibroblast and mesothelium
Myofibroblasts S-phase
Myofibroblasts G2M-phase
Myofibroblasts
Myofibroblast progenitors
Mesothelium SOX6+
Mesothelium IL18+
IP6K1
IP6K2
IP6K3
Average Expression
-1 0 1 2
Percent Expressed
0 50
Immune cells
Monocytes
Macrophages
NK and γδ cells
pre B cells
Type III ILCs
Macrophages SPP1+
Naive T cells
pDCs
DCs
Mast cells
B cells
Cell cycle
IP6K1
IP6K2
IP6K3
Average Expression
-1 0 1 2 3
Percent Expressed
0 40

**Figure S3. IP6K-expressing enteric cell subsets obtained from public scRNA-seq datasets (related to Figure 4)**

A. UMAP-based unsupervised clustering of recently reported mouse embryonic (E15.5) ENS data [S1]. Assignment of cell identities was based on the expression of signature genes—*Sox10* (Progenitor), *Ascl1* (Neuroblast), *Elavl4* (Enteric Neuron), *Plp1* (Enteric glia) and *Dhh* (SCP)—as described in the literature. Relative expression (log scale) of IP6K1 and IP6K2 among the ENS clusters.

B. Relative expression (log scale) of IP6K1 and IP6K2 in the subpopulation of human enteric cells excepting enteric neural cells, obtained by analysis of human prenatal intestinal scRNA-seq datasets [S2].

C. Relative expression (log scale) of IP6K1 and IP6K2 in mouse intestinal epithelial cells, obtained by analysis of the corresponding scRNA-seq datasets [S3]. The size and color of the dots represent the percentage of cells that express IP6K1 and IP6K2 mRNAs and their average abundances within a cluster, respectively. ENS, enteric nervous system; SCP, Schwann cell precursor; E, embryonic day; UMAP, uniform manifold approximation and projection; EEC, enteroendocrine cell; EP, enterocyte progenitor; TA, transit amplifying; CP, venous capillaries; L, large sized; M, medium sized; ILC, innate lymphoid cell; NK, natural killer; DC, dendritic cell.

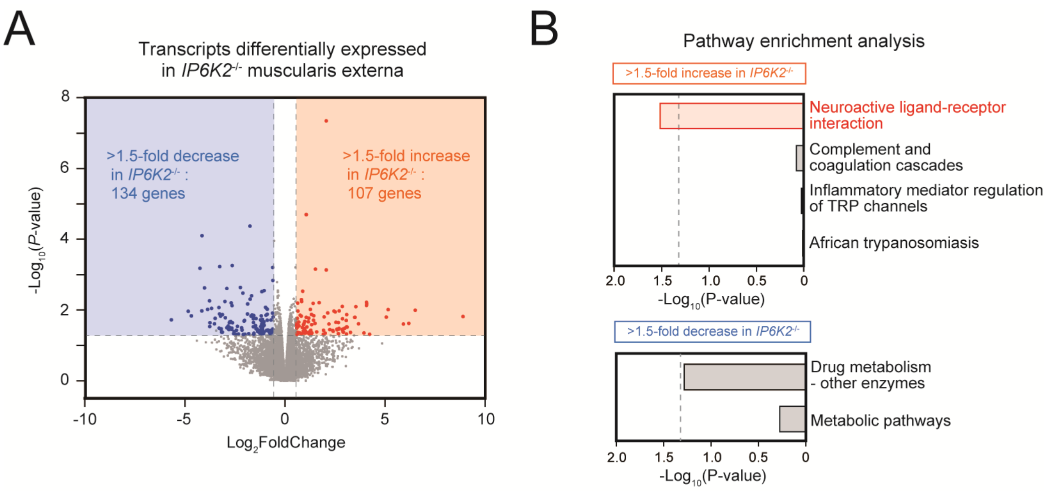


**Figure S4. RNA-seq analysis of transcripts differentially expressed in *IP6K2*$^{-/-}$ duodenal muscularis externa (related to Figure 7)**

A. Volcano plot showing transcripts differentially expressed in *IP6K2*$^{-/-}$ duodenal muscularis externa compared with WT counterparts (n = 3). Horizontal dashed line indicates P value 0.05, and vertical lines indicate 1.5-fold cutoff. Transcripts enriched or depleted more than 1.5-fold with P value < 0.05 by *IP6K2* deletion were labeled in red or blue dots, respectively.

B. Pathway enrichment analysis for transcripts with more than 1.5-fold enrichment or depletion with P value < 0.05 in the duodenum muscularis externa of *IP6K2*$^{-/-}$ mice compared with WT counterparts. KEGG pathways influenced by IP6K2 inhibition are shown. Pathways associated with neuronal regulations are highlighted in red. Vertical dashed lines indicate P value 0.05.

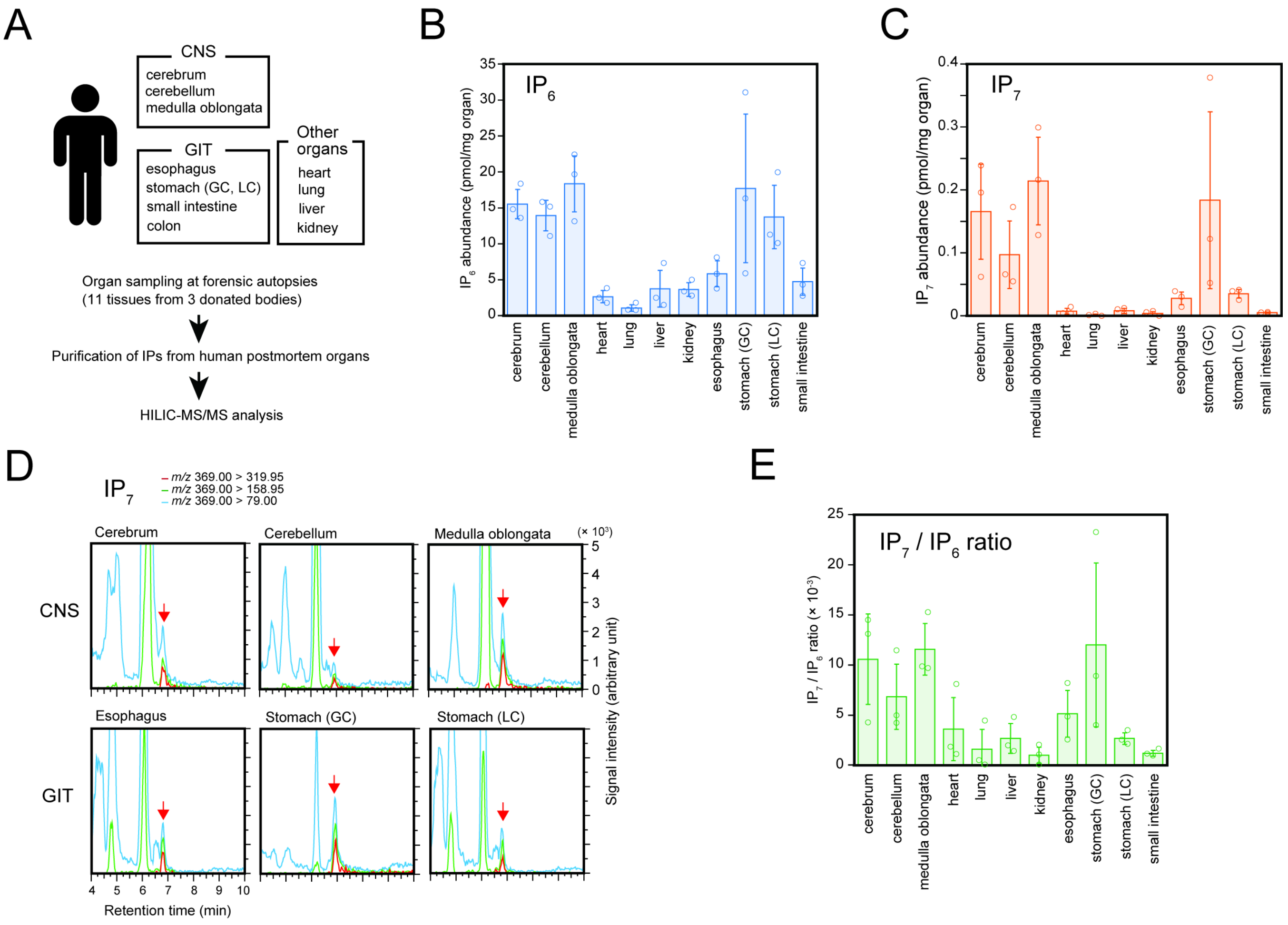


**Figure S5. HILIC-MS/MS protocol can detect $IP_7$ in human postmortem organs**

A. Schematic workflow of human organ analysis. Human postmortem organs were obtained after autopsies in three forensic cases. GC, greater curvature; LC, lesser curvature.

B, C. The concentration of $IP_6$ (B) and $IP_7$ (C) in human postmortem organs. The values shown are expressed as pmol per mg of organ weight (n = 3).

D. Representative SRM chromatograms of $IP_7$ in human postmortem CNS (cerebrum, cerebellum, and medulla oblongata) and proximal GIT (esophagus, greater curvature and lesser curvature of stomach) samples. The three best transitions are shown for $IP_7$ peak identification. Arrows indicate SRM peak of $IP_7$.

E. $IP_7/IP_6$ ratios in human postmortem organs (n = 3).

**Table S1. Optimal SRM conditions for $IP_8$ detection using HILIC-MS/MS (related to Figure 1)**

| Transition | Q1prebias (V) | Collision energy (V) | Q3prebias (V) |
|---|---|---|---|
| 818.70 > 622.95 | 22 | 33 | 20 |
| 818.70 > 720.70 | 30 | 27 | 24 |
| 818.70 > 543.10 | 40 | 45 | 40 |

**Table S2. The concentrations of $IP_6$, $IP_7$ and $IP_7/IP_6$ ratios in the 15 organs of standard diet-fed C57BL/6J mice (related to Figure 2)**

| | | cerebrum | cerebellum | spinal cord | heart | lung | liver | kidney | pancreas | spleen | gastro cnemius | testis | stomach | duodenum | small intestine | colon |
|---|---|---|---|---|---|---|---|---|---|---|---|---|---|---|---|---|
| $IP_6$ (pmol/mg organ) | #1 | 25.62 | 18.56 | 15.49 | 6.06 | 21.14 | 76.27 | 36.68 | 23.81 | 18.10 | 0.83 | 11.65 | 75.16 | 7.83 | 562.64 | 270.54 |
| | #2 | 28.87 | 16.10 | 14.37 | 5.39 | 17.21 | 45.60 | 25.19 | 26.11 | 22.36 | 1.37 | 9.69 | 132.20 | 9.26 | 148.47 | 122.52 |
| | #3 | 32.24 | 14.19 | 15.98 | 5.17 | 21.04 | 46.74 | 31.95 | 56.19 | 18.46 | 1.04 | 9.57 | 113.44 | 44.50 | 100.46 | 345.70 |
| | #4 | 17.49 | 12.48 | 14.46 | 3.23 | 10.62 | 48.83 | 29.35 | 34.01 | 16.46 | 2.44 | 9.71 | 97.19 | 8.83 | 233.85 | 34.52 |
| | average | 26.06 | 15.33 | 15.08 | 4.96 | 17.50 | 54.36 | 30.79 | 35.03 | 18.85 | 1.42 | 10.15 | 104.50 | 17.60 | 261.35 | 193.32 |
| | s.d. | 6.32 | 2.61 | 0.79 | 1.22 | 4.94 | 14.67 | 4.81 | 14.77 | 2.50 | 0.72 | 1.00 | 24.23 | 17.94 | 208.29 | 140.73 |
| $IP_7$ (pmol/mg organ) | #1 | 0.214 | 0.187 | 0.300 | 0.024 | 0.120 | 0.465 | 0.089 | 0.281 | 0.066 | 0.001 | 0.050 | 9.145 | 1.257 | 13.755 | 7.606 |
| | #2 | 0.395 | 0.288 | 0.209 | 0.035 | 0.310 | 0.206 | 0.164 | 0.560 | 0.413 | 0.000 | 0.028 | 14.751 | 1.735 | 6.724 | 2.786 |
| | #3 | 0.349 | 0.293 | 0.521 | 0.012 | 0.239 | 0.232 | 0.229 | 0.794 | 0.117 | 0.000 | 0.053 | 12.187 | 4.231 | 8.464 | 9.284 |
| | #4 | 0.267 | 0.229 | 0.350 | 0.000 | 0.085 | 0.297 | 0.061 | 0.188 | 0.071 | 0.020 | 0.047 | 8.620 | 1.480 | 12.641 | 1.797 |
| | average | 0.306 | 0.249 | 0.345 | 0.018 | 0.188 | 0.300 | 0.136 | 0.455 | 0.167 | 0.006 | 0.045 | 11.176 | 2.176 | 10.396 | 5.368 |
| | s.d. | 0.081 | 0.051 | 0.131 | 0.015 | 0.104 | 0.116 | 0.076 | 0.275 | 0.166 | 0.010 | 0.011 | 2.856 | 1.384 | 3.343 | 3.640 |
| $IP_7/IP_6$ ratio (x 1,000) | #1 | 8.34 | 10.06 | 19.34 | 3.90 | 5.67 | 6.09 | 2.43 | 11.80 | 3.65 | 1.77 | 4.31 | 121.67 | 160.56 | 24.45 | 28.11 |
| | #2 | 13.68 | 17.90 | 14.51 | 6.53 | 17.99 | 4.51 | 6.50 | 21.44 | 18.45 | 0.00 | 2.87 | 111.59 | 187.29 | 45.29 | 22.74 |
| | #3 | 10.83 | 20.63 | 32.61 | 2.30 | 11.37 | 4.96 | 7.15 | 14.12 | 6.34 | 0.44 | 5.57 | 107.43 | 95.08 | 84.25 | 26.86 |
| | #4 | 15.28 | 18.37 | 24.22 | 0.00 | 8.01 | 6.08 | 2.08 | 5.52 | 4.30 | 8.28 | 4.84 | 88.69 | 167.58 | 54.06 | 52.07 |
| | average | 12.03 | 16.74 | 22.67 | 3.18 | 10.76 | 5.41 | 4.54 | 13.22 | 8.19 | 2.63 | 4.40 | 107.34 | 152.63 | 52.01 | 32.45 |
| | s.d. | 3.07 | 4.61 | 7.72 | 2.75 | 5.36 | 0.80 | 2.66 | 6.57 | 6.94 | 3.85 | 1.14 | 13.80 | 40.00 | 24.82 | 13.28 |

**Table S3. DNA primers used for qPCR analysis in this study (related to Figure 7)**

| Gene symbol | Accession | Official name | Direction | Sequence (5' - 3') |
|---|---|---|---|---|
| Ip6k2 | NM_029634.2 | inositol hexakisphosphate kinase 2 | Forward | GAACCTGACTTCCCGCTATG |
| | | | Reverse | GTAGGATTCCTGTCGCTCCA |
| Drd5 | NM_013503.3 | dopamine receptor D5 | Forward | ACCAAGACACGGTCTTCCAC |
| | | | Reverse | CCTCCTCCTCACAGTCAAGC |
| Cckbr | NM_007627.5 | cholecystokinin B receptor | Forward | TCTCCCGCGAACTCTACCTA |
| | | | Reverse | CAGCGTTGTCATCTCCAGTC |
| Npy4r | NM_008919.4 | neuropeptide Y receptor Y4 | Forward | CCATGGCAACCTCATCTTCT |
| | | | Reverse | TCATCGATCCCTTGGATAGG |
| Nckipsd | NM_030729.4 | NCK interacting protein with SH3 domain | Forward | CCGCTGCTATCTGGAAGAAC |
| | | | Reverse | AGCACGGAAGACACCAGAGT |
| Noto | NM_001007472.2 | notochord homeobox | Forward | AATGTCACTCACCACCAGCA |
| | | | Reverse | CAGCTGGGCTCTCTCCTTC |
| Tbx1 | NM_011532.2 | T-box 1 | Forward | TGAGGAGACACGCTTCACTG |
| | | | Reverse | CTGCAGCGTCTTTGTCTGAG |
| Hrh4 | NM_153087.2 | histamine receptor H4 | Forward | AGCCTTTGTGGTGGACAGAA |
| | | | Reverse | TCGATCGTAGCTAATGAGGACA |
| Tbx18 | NM_023814.4 | T-box18 | Forward | GGATTAGACCCTCACCAGCA |
| | | | Reverse | CCTTGGTCATCCAGCTCATT |
| Pax7 | NM_011039.2 | paired box 7 | Forward | TACCAGCTGCCGGACTCTAC |
| | | | Reverse | TGACAGGGTTCATGTGGTTG |
| Mycn | NM_008709.3 | v-myc avian myelocytomatosis viral related oncogene, neuroblastoma derived | Forward | GCTGCGGTCACTAGTGTGTC |
| | | | Reverse | AAGTGGTTACCGCCTTGTTG |
| Actb | NM_007393.5 | actin, beta | Forward | CATGAAGTGTGACGTTGACATC |
| | | | Reverse | ATGATCTTGATCTTCATGGTGC |
| Rn18S | NR_003278.3 | 18S ribosomal RNA | Forward | GTAACCCGTTGAACCCCATT |
| | | | Reverse | AGTTCGACCGTCTTCTCAGC |
